# Uncovering bi-directional causal relationships between plasma proteins and psychiatric disorders: A proteome-wide study and directed network analysis

**DOI:** 10.1101/648113

**Authors:** Carlos Kwan-long Chau, Alexandria Lau, Pak-Chung Sham, Hon-Cheong So

## Abstract

Psychiatric disorders represent a major public health burden yet their etiologies remain poorly understood, and treatment advances are limited. In addition, there are no reliable biomarkers for diagnosis or progress monitoring.

Here we performed a proteome-wide causal association study covering 3522 plasma proteins and 24 psychiatric traits or disorders, based on large-scale GWAS data and the principle of Mendelian randomization (MR). We have conducted ~95,000 MR analyses in total; to our knowledge, this is the most comprehensive study on the causal relationship between plasma proteins and psychiatric traits.

The analysis was *bi*-directional: we studied how proteins may affect psychiatric disorder risks, but also looked into how psychiatric traits/disorders may be causal risk factors for changes in protein levels. We also performed a variety of additional analysis to prioritize protein-disease associations, including HEIDI test for distinguishing functional association from linkage, analysis restricted to *cis*- acting variants and replications in independent datasets from the UK Biobank. Based on the MR results, we constructed directed networks linking proteins, drugs and different psychiatric traits, hence shedding light on their complex relationships and drug repositioning opportunities. Interestingly, many top proteins were related to inflammation or immune functioning. The full results were also made available online in searchable databases.

In conclusion, identifying proteins causal to disease development have important implications on drug discovery or repurposing. Findings from this study may also guide the development of blood-based biomarkers for the prediction or diagnosis of psychiatric disorders, as well as assessment of disease progression or recovery.

## Introduction

Psychiatric disorders as a whole represent a major public health burden and are leading causes of disability worldwide^1^. Nevertheless, there remain significant challenges in the diagnosis and treatment of psychiatric disorders. The etiologies of most psychiatric disorders are not fully understood yet, and we still have limited knowledge about the factors underlying disease progression or recovery. There have been limited advances on the treatment of psychiatric disorders, especially on the discovery of drugs with novel mechanisms of actions.

Another longstanding challenge is that unlike many other medical diseases, there are yet *no* reliable biomarkers for diagnosis, prognosis or monitoring progress of psychiatric disorders^2^. The biomarker should preferably easily accessible and measured, and blood/plasma proteins represent one type of biomarkers fulfilling such requirement. A number of studies have investigated into differences in protein levels in various psychiatric disorders, many of which are hypothesis-driven^2, 3^. The rise of high-throughput proteomic platforms now enables interrogation of a large panel of proteins at the same time. However, as pointed out by a recent review, there were several important limitations in these studies, for example inadequate sample sizes, methodological heterogeneity, and risk of confounding^4^.

An important question underlying many protein marker (or proteomics) study in observational samples is the difficulty in assessing causality and the direction of effect. Is the change in the level of a protein the cause or consequence of the disorder? Ideally one may follow-up a group of subjects at risk for the disorder and regularly monitor the protein level to ascertain the temporal relationship between disease onset and rise or fall in protein levels. However, this is often impractical due to low incidence of some disorders, the cost and difficulty in identifying at-risk subjects, and loss to follow-up. Another important concern is confounding bias. For example, patients are often medicated at the time of recruitment, and numerous other known or unknown confounders (e.g. age, gender, stressors, comorbid medical conditions) can result in spurious associations between a protein and the psychiatric disorder.

The recent decade has witnessed a rapid growth in genomic research in psychiatric disorders; in particular, high-throughput technologies such as genome-wide association studies (GWAS) have been highly successful in unraveling genetic variants associated with many neuropsychiatric traits. With the advent of GWAS, a new approach to causal inference, known as Mendelian randomization (MR), is gaining increased attention. An MR study can be considered a “naturalistic” randomized controlled trial (RCT)^5, 6^ The basic principle is to use genetic variants as “instruments” to represent the actual risk factor, and analyze its relationship with the outcome. For example, a person who has inherited a low-density lipoprotein (LDL) lowering allele (or allelic score) at a relevant genetic locus is analogous to receiving a LDL-lowering drug, and vice versa. To estimate the causal effects of reduced LDL, we can then test whether people who have inherited the lipid-lowering genetic variants are less likely to develop cardiovascular diseases. The random allocation of alleles at conception is analogous to randomization in clinical trials; as such, MR studies are less vulnerable to confounding bias than observational studies^5^. And since a person’s genetic profile is fixed at birth, this approach also avoids problems of reversed causality.

In this work we aim to investigate potential casual relationships between plasma protein levels and a range of psychiatric disorders/traits, based on the Mendelian randomization approach. As discussed above, a major advantage is that MR is less susceptible to confounder bias and reversed causality. MR also enables us to test direction of causal effects in both directions (protein-to-disease as well as disease-to-protein). In addition, MR can be performed using GWAS summary statistics, which are usually based on very large samples (usually >10000). On the contrary, the sample sizes of previous proteomics studies are typically small (*N* mostly <100; see ref^4^). The MR approach also provides a cost-effective way to explore a wide panel of proteins against a variety of psychiatric traits, which will be prohibitively costly to carry out in a conventional longitudinal or observational study. Here we made use of recently published results of proteomic quantitative trait loci (pQTLs) and the latest large-scale GWAS data from the Psychiatric Genomics Consortium (PGC), UK Biobank and other sources to perform MR analysis.

While many of the MR studies were hypothesis-driven and focused on testing causal associations of known risk factors, MR can also be extended to screen for a larger number of risk factors to uncover novel causal associations. In the field of psychiatric genetics, hypothesis-driven candidate gene studies have now turned out to be largely irreproducible^7, 8^, suggesting that candidate-based approaches to ‘omics’ studies have important limitations. Here we adopted a systematic and unbiased ‘hypothesis-generating’ or ‘hypothesis-free’ approach and scans all available proteomic markers for casual relationships with psychiatric disorders in both directions.

By exploring proteins causal to the development of psychiatric disorders, we will gain a better understanding into the pathogenesis of psychiatric disorders. Of note, many susceptibility SNPs found in GWAS reside in non-coding or inter-genic regions, and their functional impact remains to be elucidated^9^. Linking SNPs to protein QTLs will help to reveal the functional and biological impact of susceptibility variants. From a clinical point of view, drugs that target the causal proteins may be effective for disease prevention and treatment. If such drugs are already in use, they can be repositioned for the disorders, saving the high cost and time for new drug development. In addition, proteins with causal relationship with the disease may also serve as predictive biomarkers for at-risk patients before full development of the disorder.

On the other hand, the study of proteins whose levels are affected *as a consequence* of disease liability also has important implications. The discovery of such proteins may enhance our understanding of the molecular mechanisms underlying disease progression and complications. From a clinical perspective, such proteins may be candidate biomarkers for monitoring and assessing disease progress or recovery. An important definition of causality is that intervention on the cause will lead to changes in the outcome. If a disease is a *causal* risk factor for protein level change (let’s say increase), then when the disease is cured, the protein level will change (decrease) accordingly. The reverse (i.e. rise in protein level) may be observed when the disease becomes more severe or during relapses. This may be particularly relevant in psychiatry, as patient progress is usually monitored by clinical observations and self-reported symptoms, with no objective biomarkers available.

In addition to the above, sometimes a protein whose level changes as a result of a disease may yet be a risk factor for another condition. For example, depression is associated with increased risk of cardiometabolic disorders^10^, and some have postulated inflammatory pathways and proteins as mediating factors^11, 12^. We may decipher mediating proteins between different psychiatric disorders/traits by studying bi-directional causal links between proteins and diseases. In this study, we have presented a framework for *directed network* (a network in which edges have directions) analysis of psychiatric traits and proteins based on our bi-directional MR results, and external protein-protein interaction (PPI) databases.

As a summary, in this work we first employed two-sample MR to identify proteins causally associated with psychiatric disorders/traits, with replication in the UK Biobank sample. Next we conducted MR analysis with psychiatric disorders/traits as exposure and protein levels as outcome. These analyses were conducted in a large-scale and systematic manner covering up to 3522 plasma proteins and 24 GWAS datasets of psychiatric disorders/traits. Based on these results, we constructed directed protein-disease networks which aimed to shed light on the pathways between various proteins and psychiatric disorders. The full analysis results are made publicly available, which we believe will serve as a useful resource for further clinical or experimental studies. An overview of our analytic framework is presented in Figure 1.

**Figure 1.**
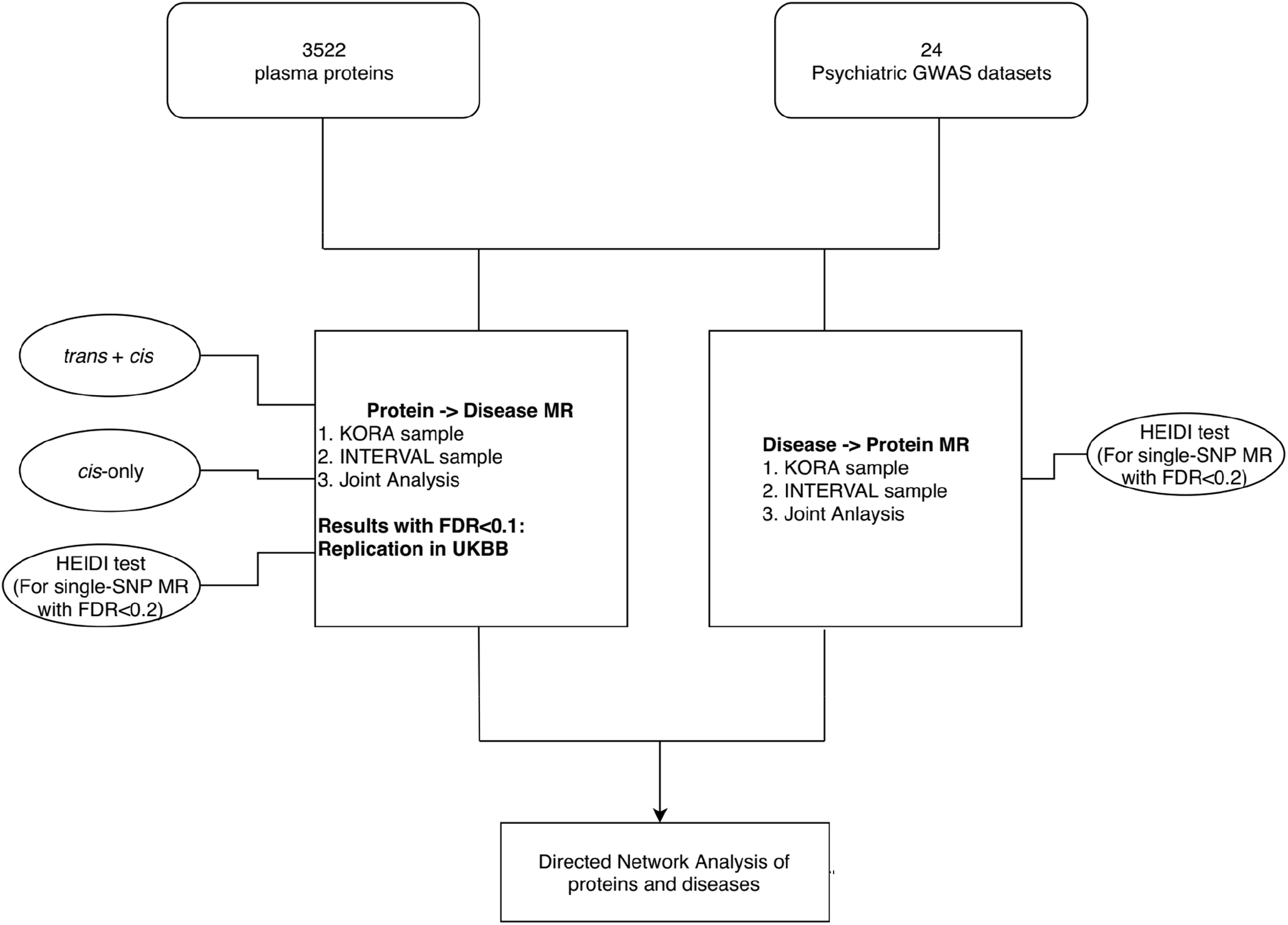
An overview of the analytic framework. Briefly, we conducted MR analysis with psychiatric disorders/traits as exposure and protein levels as outcome. The analysis was conducted in a large-scale and systematic manner covering up to 3522 plasma proteins and 24 GWAS datasets of psychiatric disorders/traits. We also conducted several additional analyses, including a cis-only analysis (as cis-variants are as instruments less likely to be affected by pleiotropy), HEIDI test for selected associations to distinguish ‘linkage’ from functional association, as well as replication in UK Biobank of top protein-to-disease associations. Based on these results, we constructed directed protein-disease networks which aimed to shed light on the pathways between various proteins and psychiatric disorders.

## Methods

### Blood Plasma Protein pQTL Datasets

Blood plasma protein pQTL data were taken from the recently published KORA and INTERVAL studies^13, 14^.

Full summary statistics of the two proteomics GWAS are downloaded from https://metabolomics.helmholtz-muenchen.de/pgwas/index.php?task=download and https://www.phpc.cam.ac.uk/ceu/proteins/. KORA is a German cohort with GWAS performed on levels of 1124 protein levels from a population-based cohort of 1000 subjects. Proteins were quantified by an aptamer-based affinity proteomics platform known as SOMAscan. While the original paper by Suhre et al.^13^ also included a sample from the QMDiab study, that sample was recruited from Arab and South Asian countries, and was excluded from our analysis since most other GWAS datasets we analyzed were from Europeans. The INTERVAL study is a similar proteomics study from a UK cohort, investigating the association of 3622 plasma proteins in 3301 healthy subjects^14^. Again the SOMAscan platform was employed for protein quantification. For details of each dataset, please refer to the original papers^13, 14^.

We have included these two studies based on their relatively large sample sizes, the use of same platform for protein quantification, and the fact that both studies have full GWAS summary statistics available for download. Full proteomics GWAS data are required for our MR analysis to identify which proteins may be affected as a consequence of liability to psychiatric disorders and for HEIDI analysis.

### Psychiatric GWAS datasets

We extracted psychiatric GWAS summary datasets mainly from the Psychiatric Genomic Consortium (PGC) and UK Biobank (UKBB). We included in total 24 GWAS datasets, which encompass a comprehensive range of psychiatric disorders (and traits closely linked to the disorders). They include attention deficit and hyperactivity disorder (ADHD)^15^, anorexia nervosa (AN)^16^, Alzheimer’s disease (ALZ)^17^, autistic spectrum disorders (ASD)^18^, anxiety disorders^19^ (including generalized anxiety disorder, phobic disorder, social phobia, agoraphobia, and specific phobias; results from both case-control study and derived factor score included), depressive symptoms (DS)^20^, major depressive disorder (MDD)^21, 22^ (two sources, one sample^22^ comprised female patients enriched for severe melancholic depression), longest period of depressed mood, number of episodes of depressed mood (GWAS from UK-Biobank data from Neale’s lab^23^), neuroticism (NEU)^20^, obsessive compulsive disorder (OCD)^24^, post-traumatic stress disorder (PTSD)^25^, bipolar disorder (BIP)^26^, schizophrenia (SCZ)^27^ and history of deliberate self-harm (DSH) or suicide^23^. We also include GWAS on antidepressant response (as both binary and quantitative outcome)^28^, and a GWAS on SCZ versus bipolar disorder patients^29^. For ADHD and PTSD, we also performed separate analysis on male-only, female-only and combined samples, as the corresponding summary statistics has been made available and there is good evidence for gender differences in these disorders^30, 31^.

For some of the psychiatric disorders/traits, a large-scale (‘primary’) study is available from PGC or a related source, while a corresponding trait is also recorded in UKBB. These psychiatric disorders/traits include anxiety disorders, ASD, BIP, OCD, PTSD, AN, SCZ and ALZ (parent’s ALZ disease status was used as a proxy for analysis). The details are given in Table S3. For these disorders/traits, we also included the UKBB association dataset as a replication set when we studied the causal effects of proteins on diseases.

### MR analysis: protein as exposure and psychiatric disorder/trait as outcome

We first treated plasma protein levels as exposure and psychiatric disorders/traits as outcome. As discussed above, for some of the disorders/traits, a large-scale study dedicated to the study of the specific disorder is available in PGC (or a similar source such as IGAP^17^), while a similar corresponding trait was also recorded in the UKBB sample. In such cases, we would conduct a ‘primary’ analysis based on the PGC data, then extract the top results (FDR<0.1, based on combined *cis & trans* pQTLs) for replication analyses using the UKBB data.

We extracted pQTL according to those provided in the original papers (Supplementary data 1 in Suhre et al.^13^ and Supplementary Table 4 in Sun et al.^14^). Exposure instrument SNPs were harmonized with psychiatric GWAS summary data with the harmonise_data function in the R package “TwoSampleMR”. LD-clumping was performed with the clump_data function in “TwoSampleMR” at an r-squared threshold of 0.1 (within 10000 kb). We set a more liberal r-squared threshold than the default (0.001) to allow possibly more instruments to be included. The threshold was set to a relatively low level such that only weak LD is allowed, in order to prevent instability in MR estimates. Residual SNP correlations were properly taken into account of in our inverse-variance weighted (IVW) and MR-Egger analyses (see Supp. Text)^32, 33^.

#### Statistical analysis methodology

The primary method of MR analysis is based on the number of SNPs (Table 1). If there is evidence of significant pleiotropy as revealed by a significant intercept term in Egger regression, we would employ the MR-Egger approach, which enables valid causal estimates to be given in the presence of imbalanced horizontal pleiotropy. Here horizontal pleiotropy refers to instrument SNPs having effect on the outcome through pathways other than the exposure. Otherwise, the standard Wald ratio or IVW approach was used. The R package “MendelianRandomization” was employed, which allowed correlations between instrument SNPs to be accounted for.

**Table 1.** MR approach adopted according to the number of instrument SNPs

| Number of SNP in<br>Harmonized Data (*n*) | 2MR Test Applied |
| --- | --- |
| $n = 1$ | Wald Ratio |
| $n = 2$ | Inverse variance weighted (IVW) |
| $n \geq 3$ | If (pleiotropy *p*-value < 0.05)<br>Egger's Regression<br>else<br>Inverse variance weighted |

To control for multiple testing, we employed the false discovery rate (FDR) approach, which controls the expected proportion of false positives within the rejected hypotheses. FDR was computed by the Benjamini-Hochberg method^34^ and a modified approach by Storey^35^. The latter approach (implemented in the R package “qvalue”) also estimates the proportion of null which could lead to improvement in power (this is the default method used in this study). Note that the FDR approach is also valid under positive (regression) dependency of hypothesis tests^36^. The FDR-adjusted *p*-value, or *q*-value, refers to the lowest FDR incurred when calling a feature significant^35^.

#### Other sensitivity tests

##### Analysis restricted to *cis*-acting pQTLs

Both studies classified pQTL-plasma-protein association as *cis* or *trans*, and in this work we would perform analysis on *cis + trans* pQTLs combined and *cis*-pQTLs only.

We primarily present results from analysis of combined *cis*- and *trans*-pQTLs, as this would allow more instruments to be included, improving the power and reliability of results, and would enable methods such as MR-Egger to be performed to correct for imbalanced horizontal pleiotropy. We noted that a substantial proportion of pQTLs identified by Suhre and Sun et al. and were in fact *trans*-pQTLs (~27% and 70% respectively), hence discarding *trans*-acting variants will result in substantial waste of available data and loss of statistical power.

Nevertheless, c*is*-acting pQTL have strong prior probability of association and are less susceptible to horizontal pleiotropy, although they can also affect the outcome via other pathways. Here we also performed additional analysis by sub-setting the original data to produce 2 extra exposure datasets (named *cis*-KORA, *cis*-INTERVAL), and MR was repeated with *cis*-pQTLs only.

##### HEIDI test

Sometimes the instrument SNP can be in LD with another SNP, which is causally associated with the outcome. This scenario is known as ‘linkage’ by Zhu et al.^37^, which may be distinguished with functional associations (same SNP affecting both protein level and outcome) by the SMR-HEIDI test^37^. The test was designed especially for single-SNP MR analysis with the assumption of a single causal variant in the region. We employed the SMR program (https://cnsgenomics.com/software/smr/#SMR) written by the authors. Default settings were used, in which up to the top 20 pQTLs in the *cis* region were extracted for the HEIDI test. The test requires summary statistics of the proteomics GWAS, and is relatively computationally time-consuming to perform. We conducted HEIDI for single-SNP MR results with FDR<0.2 to limit the total number of tests to several hundreds. An HEIDI *p*-value >0.05 is marked as ‘passing’ the test as per the original authors, but due to multiple testing of a large number of hypotheses, we also presented FDR-adjusted *p*-values.

In practice, we note that occasionally HEIDI would fail to produce results, presumably due to inadequate number of SNPs in the *cis* region of target pQTL passing the preset *p*-value threshold (1.57E-3, equivalent to chi-square statistic of 10), or some variants may be filtered away due to too low or too high LD. Another important limitation is that currently HEIDI or other co-localization methods cannot produce reliable results with the number of causal variants >=2. For these reasons we included HEIDI as a sensitivity test, but we do not exclude associations failing this test from subsequent analyses. If multiple testing is accounted for by the FDR approach, a relatively small proportion of HEIDI results remained significant after FDR correction (see Results).

#### Replication Analysis

We sought replication for the results from our primary analysis (mainly PGC studies) in an independent sample from the UK Biobank. Replication was performed for three sets of MR results (each result represents a protein-disorder pair): (1) MR results based on KORA sample with FDR<0.1; (2) Results based on INTERVAL sample with FDR<0.1; (3) Results from joint KORA + INTERVAL meta-analysis with estimated FDR<0.1 (based on Simes *p*-value). In (3) only overlapping proteins in the two samples were included.

As for the methodology of joint analysis, we computed the maximum *p*-value (max_*p*) as a test of the hypothesis of non-null associations in *both* samples, as described by Nichols et al^38^. On the other hand, the Simes test^39^ was conducted to combine the *p*-values to test the global null hypothesis (i.e. H_1_: association in *any* sample). The max*_p* and Simes test procedures are both robust to (positive) dependencies of hypothesis tests^38, 39^. We also computed a standard IVW meta-analysis with “metagen” in R to obtain an aggregate estimate of causal effects from both PGC studies and UKBB. (Strictly speaking the statistical significance from a standard IVW meta-analysis may be inflated due to dependency of effect estimates, but this was included as an additional approach to prioritizing associations and for computing the overall effect sizes). We adopted a one-sided test for computing *p*-value in the replication sample.

We also attempted a more novel analysis known as replicability FDR (repFDR), which assesses the expected proportion of false replications^40^. This is a stringent test for association in every sample. H_1_ (alternative hypothesis) refers to non-null results or true associations in *every* testing sample, taking into account the number of hypothesis tested. Intuitively, when a large number of hypothesis is tested, to achieve low repFDR the hypothesis needs to be highly significant (having low FDR) in every sample. The authors proposed the “*r* value”, which is defined as “the lowest FDR level at which we can say that the finding is among the replicated ones”^40^. The analysis was carried out with default settings using an online program provided by the authors (http://www.math.tau.ac.il/~ruheller/App.html).

### MR in the other direction: Psychiatric disorders/traits as exposure and proteins as outcome

We then performed MR analysis in the other direction in which the 24 psychiatric disorders/traits were treated as exposure, while plasma protein levels were considered as outcome.

SNPs in psychiatric GWAS datasets were first selected based on the genome-wide significance threshold (p < 5E-08), then clumped at r^2^=0.1 (within 10000 kb). The rest of the analysis follows what was described in the earlier sections. The HEIDI test was performed for single-SNP analyses. In this case we have two independent outcome datasets (KORA and INTERVAL), so we also performed replication of the results from the first dataset on the second one. We conducted meta-analysis on the intersecting proteins and repFDR was computed as described above.

#### Conversion of coefficients from UKBB summary data

The UKBB GWAS results are based on linear regression of the outcome, regardless of whether it is binary or quantitative (details available at https://github.com/Nealelab/UK_Biobank_GWAS). For consistency and ease of meta-analysis with other GWAS datasets based on logistic regression, we converted the linear regression coefficients to those under a logistic model, based on Lloyd-Jones et al.^41^. Standard error of the converted coefficients was derived by the delta method.

### Directed network of proteins, drugs and psychiatric traits/disorders

Here we constructed a directed network to link up different proteins and psychiatric traits/disorders for better visualization of their relationships; drugs targeting proteins identified in MR were also included in the network. We restricted our attention to intersecting proteins in the KORA and INTERVAL datasets. To be included in the network, the exposure-outcome pair should have an FDR <=0.1 based on joint analysis. The edge weights reflect the strength of association between the disease and protein, derived from the average regression coefficients from MR using the KORA and INTERVAL samples. The above criteria were chosen to balance interpretability and comprehensiveness of the network. More formally, we visualized the causal relationships between protein and diseases as a network G(V,E), where V are unique elements from the set *S* = {(284 KORA proteins ∩ 1487 INTERVAL protein) ∪ 24 psychiatric disorders} and E (edges) = {(p_1_,d_1_),(p_2_,d_1_),…,(p_1_,d_2_),(p_2_,d_2_),…,(p_n_,d_k_),…,(d_i_,p_n_)}, where p_n_ indicates protein vertices and d_k_ and d_i_ indicate vertices of psychiatric disorders/traits.

We also extracted drugs that target proteins identified in our MR analysis based on DGIdb (ver 3.0;^42^) to highlight drug repositioning opportunities.

The final outcome is a network with 151 nodes and 236 edges. The graph is visualized with the R package RCyp3 (version 2.4.6) and Cytoscape (version 3.7.2).

### Literature-pruned protein-disease network

To prioritize known protein-disease or disease-protein pairs that are strongly supported by literature evidence, we perform a systematic PubMed search for each of the unique 213 pairs. Queries are entered as “(EXPOSURE) AND (OUTCOME)” using the R package “easyPubMed” (version 2.13). We set the maximum results returned to be 20. Results are also visualized with a graph similar to the above network, but the edge weights are given by the number of matches returned by PubMed. We make no assumptions about the directions of the associations thus no direction is indicated.

### Pathway enrichment analysis of the proteins in the directed network

To examine the global functional significance of our MR results of blood proteomics and psychiatric disorders, we performed a graph-wide functional annotation using ClueGO (version 2.5.5). Gene-sets and pathways were extracted from ClinVar, KEGG, Reactome Pathways, Reactome Reactions and WikiPathways. Enrichment significance was estimated with hypergeometric tests and multiple comparisons were accounted for by Bonferroni correction. A network was constructed to visualize the relationship between the pathways and proteins. The size of each node correlates with the level of significance.

### Online searchable databases

To facilitate searching of results, we also constructed *online databases* for Tables S1 to S4. Please refer to ‘Availability of Supplementary Materials’ for details.

## Results

### MR analysis: Proteins as exposure and psychiatric disorders as outcome

#### Main analysis based on the KORA and INTERVAL studies

The pQTLs of KORA and INTERVAL studies are listed in Tables S1a and S1b. In total 539 pQTLs were extracted from the KORA sample, of which 391 are *cis* variants. In total 1980 pQTLs were extracted from the INTERVAL data, of which 589 are *cis* variants. The KORA and INTERVAL datasets covered 284 and 1478 proteins respectively.

We have conducted totally 6038 and 25034 MR tests based on the KORA and INTERVAL samples respectively. Each test examined the effect of a protein on the psychiatric disorder/trait under study. We also repeated the analysis restricted to *cis* variants only, with 4285 and 8201 MR tests performed on KORA and INTERVAL respectively.

The main MR analysis results are presented in Table 2 and Supplementary Tables S1c to S1d. For the analysis based on KORA pQTLs, 13 protein-disease pairs were significant at an FDR level of 0.1 and 20 at FDR<0.2. For the analysis using pQTLs from INTERVAL sample, we found 391 associations at FDR<0.1 and 456 at FDR<0.2.

**Table 2.**
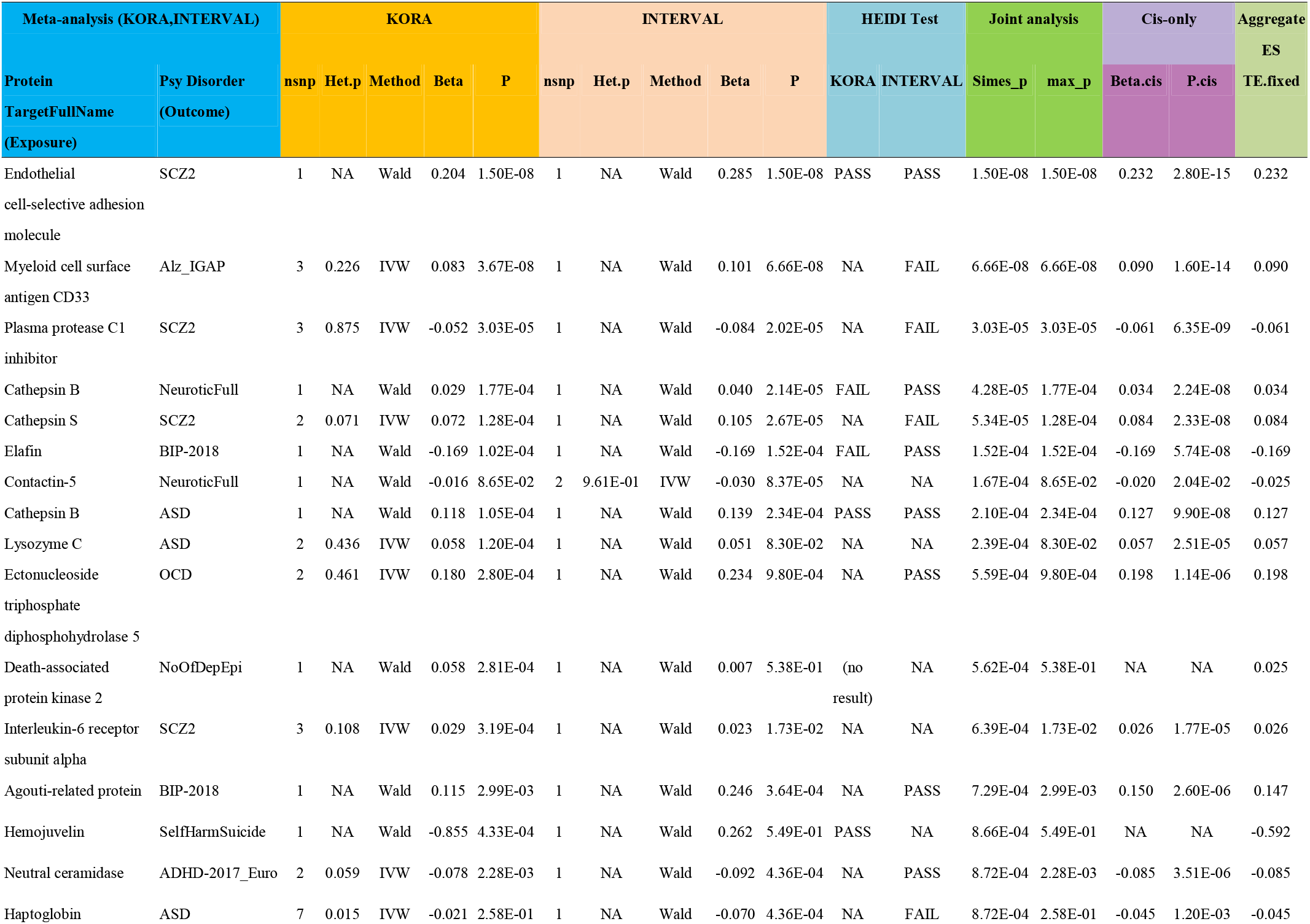
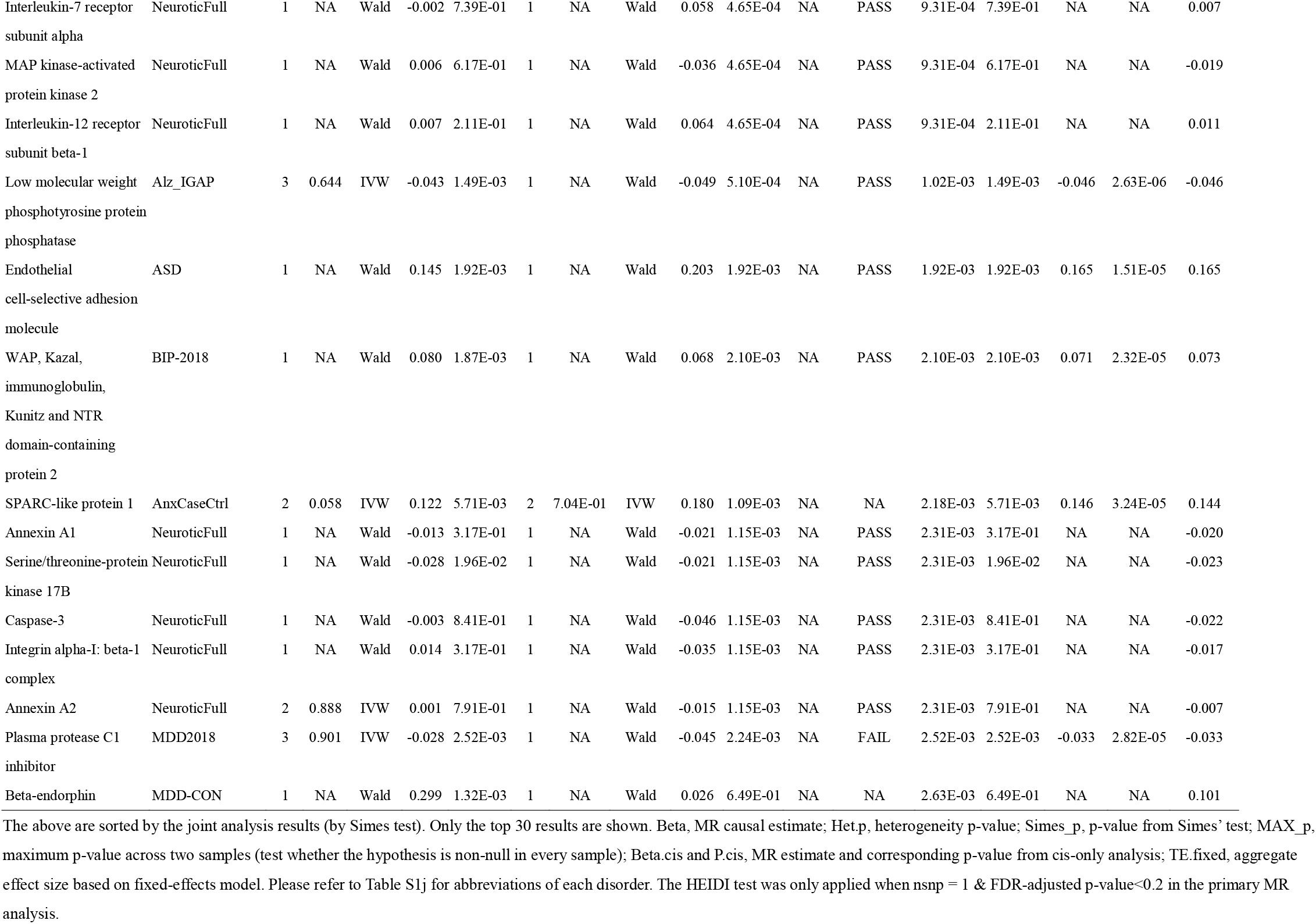
MR results with plasma proteins as exposure and psychiatric disorders/traits as outcome (joint analysis of KORA and INTERVAL; top 30 results shown)

#### HEIDI test

We also performed HEIDI test as a sensitivity test to detect associations that may be due to linkage (Table S1c, S1d). The original authors used the conventional threshold *p*<0.05 as the threshold for passing or failing the test; however, we also note there is a high chance of false positives in view of the large number of tests performed. We computed FDR with the Benjamini-Hochberg method; among the protein-disease associations tested by HEIDI, a relatively modest proportion have low FDR-adjusted *p*-values<0.05 (overall 4.1 %), <0.1 (5.2%) or <0.2 (6.2%). It should be noted that failing the HEIDI does not always mean the result is invalid. For example, there can be other true causal variant(s) affecting the protein and outcome at the same time, which is/are also in LD with the target pQTL. However, the test is still useful in prioritizing proteins for further studies.

#### Joint analysis

We also performed a joint analysis on the intersecting proteins between the two pQTL datasets (Table 2 and Table S1e) to estimate of the aggregate causal effects. The results of IVW meta-analysis, maximum *p* (a test for non-null association in every sample) and Simes test (a test for non-null association in at least one sample) are given in Table S1e. Top 30 results sorted by Simes *p*-value are presented in Table 2.

We also performed ***cis*-only** analysis and the results are summarized in Table S2. The *cis*-pQTLs are summarized in Table S2a and S2b for reference. Results of KORA- or INTERVAL-only analysis are given in Table S2c and S2d, while joint analysis is presented in Table S2e. In total 12 and 15 associations achieved FDR<0.1 and <0.2 in the KORA sample respectively; 21 and 32 associations achieved FDR<0.1 and <0.2 in the INTERVAL sample respectively.

#### Cis-only analysis

There is significant overlap in the results between the *cis*-only and combined *cis & trans* pQTL analyses. For example, 68 and 45 protein-disease associations showed Simes *p*< 0.01 in the *cis & trans* and *cis*-only analysis respectively, among which 44 of them overlapped (Table S1e). Note that we also provide a summary of the *cis*-only analysis results alongside the main results (*cis + trans*) for easy reference.

#### Highlights of top protein-to-disease associations

We found a number of interesting protein-disease associations among the top results, and shall highlight a few here. Here our discussion is restricted to the top ~50 results in the joint analysis, as listed in Table S1e. For instance, cathepsin B is found to be associated with increased ASD risk. Interestingly, a recent study reported that inhibition of cathespin B led to a reverse of increased leukocyte-endothelial adhesion (which is implicated in neuro-inflammation) in the cerebral vessels in an autistic mouse model^43^. As another example, we found that interleulin-6 receptor subunit alpha is positively associated with SCZ. Inflammation is believed to play an important role in the pathogenesis of SCZ^44^, and clinical studies have also found rise in inflammatory markers in SCZ^44^. Soluble IL-6 receptor (sIL-6R) may be involved in ‘trans-signaling’, in which the complex of IL-6 and sIL-6R may bind to gp130 and exert pro-inflammatory effects^45^. Another recent MR study also reported increased sIL-6R being associated with SCZ^46^, but here the proteomics dataset used is different and the association is more specific (for the alpha subunit). The exact role of IL-6 pathways in SCZ remains to be elucidated in future studies. Also in line with the inflammation hypothesis in SCZ and other psychiatric disorders (e.g. ASD^47^), we observed that the protein endothelial cell-selective adhesion molecule (ESAM) was positively associated with SCZ and ASD risks. Neurovascular endothelial dysfunction as well as hyper-permeability of the blood brain barrier (BBB) may be linked to neuroinflammation, which has been implicated in a number of neuropsychiatric disorders^48, 49^.

Another interesting finding is that increased beta-endorphin levels were found to be causally associated with elevated risk of melancholic depression (the MDD-converge sample mainly recruited subjects with melancholic depression). Melancholic depression has been reported to be associated with hypothalamic-adrenal (HPA) axis hyper-activation, while beta-endorphin may be involved in the stimulation of HPA axis^50, 51^. In clinical studies, both increase and decrease in beta-endorphin levels have been reported; yet some studies specifically focused on melancholic patients indeed reported elevation of beta-endorphin levels (please refer to Table 1 of Hegadoren et al.^52^). These findings call for the need of more homogeneous subgrouping of MDD patients in biomarker studies.

Yet another finding that may be of clinical relevance was that lower ferritin was found to be casually linked to increased ASD risks. Reduced ferritin and iron deficiency were reported to be common among children with ASD^53, 54^. Interestingly, a clinical study reported that iron supplementation improves sleep disturbance in ASD patients^55^, although this is a case-only study without a placebo control arm.

Taken together, our findings help shed light on the pathogenesis of psychiatric disorders. In certain occasions if drugs are available to target the tentative causal proteins, such drugs may be repurposed to treat the corresponding disorders.

#### Replication analysis

Replication analysis was performed for MR results having FDR<0.1 in the KORA sample, INTERVAL sample or in joint analysis (Table S3). For the KORA dataset, altogether 6 associations were replicated with one-sided *p*-value <0.05 in the UKBB dataset (spanning three unique proteins) (Table S3b). Note that sometimes more than one UKBB dataset may correspond to the original phenotype and we included all of them in the replication analysis. The use of parental history of Alzheimer disease as a proxy of the disease phenotype has been successfully adopted in previous works^56^ and this strategy was also employed in our replication analysis. In the INTERVAL sample, 97 associations achieved replication *p*-value<0.05 in UKBB (spanning 51 unique proteins), among which 77 had replication FDR<0.05 (spanning 46 unique proteins) [Table S3c].

We observed that many of the top results were from Alz-IGAP or SCZ; this is likely due to relatively large sample size and power of these two datasets. However, some of the top results for Alzheimer disease showed heterogeneity of instrument effects. This can be due to various reasons^57^, but horizontal pleiotropy could be one, therefore such results might be viewed with caution. For comprehensiveness, we also extracted a set of intersecting proteins between the two pQTL datasets with FDR<0.1 in joint analysis (by Simes test), and carried out meta-analysis combining PGC results with UKBB, using either KORA (Table S3d) or INTERVAL (Table S3e) pQTL data.

We briefly highlight a few replicated protein-to-disease associations (we only focus on those listed in Table S3d/e due to the large number of associations). For example, protease C1 inhibitor, a protein involved in immune functioning, was among the top replicated associations for SCZ. Inflammation and immune pathways have been strongly implicated in the etiology of SCZ^44^. Protease C1 inhibitor an inhibitor of the classical pathway of complement and the contact system^58^, which was reported to reduce the release of various cytokines in a sepsis model of primates. We observed that reduced levels of C1 inhibitor was causally linked to SCZ. This suggests that increased inflammatory responses, at least partially mediated by the complement pathways, may be a causal risk factor in the pathogenesis of SCZ. Myeloid cell surface antigen (CD33) is another replicated causal association with Alzheimer disease (Table S3d/e), which has been reported to be associated with increased neuritic amyloid pathology and microglial activation^59^, and has been suggested as a promising therapeutic target^60^.

#### Relationship to drug repositioning or discovery

If a protein is causal to a psychiatric disorder, then a drug which targets the protein as an antagonist or inhibitor may be able to treat the disease if increased protein levels are casually associated with the disease. On the contrary, the psychiatric disorder may be a side-effect of that drug if reduced protein levels are associated with disease. The reverse is true for the drugs acting as agonists. To facilitate the discovery of candidates for drug repositioning and the exploration of medication side-effects, we also searched ChEMBL and DrugBank database for existing drugs for top proteins in joint analysis (Table S1f/S5b)

We will highlight and discuss several promising repositioning opportunities based on proteins with larger effect size on diseases, and those that can be targeted by existing drugs.

##### NAAA and psychiatric disorders

We found that high N-acylethanolamine-hydrolyzing acid amidase (NAAA) is causally associated with higher AD risk (KORA *β*=0.038, INTERVAL *β*=0.062, Simes P=0.04) and PTSD risk (KORA *β*=0.21, INTERVAL *β*=0.21, simes P=0.02). NAAA potentially affects the pathophysiology of psychiatric disorders with its role in the hydrolysis of N-acylethanolamines (NAEs). NAEs are structural analogues to the endocannabinoid arachidonoylethanolamide (anandamide), and their functions are beginning to be elucidated. NAEs may be involved in lipid metabolism and have anti-inflammatory effects^61^. Besides, NAEs may also modulate brain functions and hence implicated in neuropsychiatric disorders, through their effects on cognitive functions, mood, reward and sleep-wake cycles etc.^62^.

NAAA is involved in the metabolism of endocannabinoids^63, 64^, which may play a role in PTSD and AD^65-68^. Studies of endocannabinoids as a biomarker in PTSD also showed promising results^69^ In fact, synthetic endocannabinoid (nabilone) has been shown to be effective in relieving symptoms of PTSD for the majority of patients (~72%) in an open label clinical trial^70^. Thus, NAAA inhibitors may be drug candidates for PTSD, AD or other psychiatric disorders. A number of NAAA inhibitors have been developed^71^. Apart from the above, NAAA is also highly effective in palmitoylethanolamide (PEA) degradation, while PEA has been reported as a neuroprotective agent and therapeutic target for AD^72^.

##### IL-6 signaling and psychiatric disorders

We also found that interleulin-6 receptor subunit alpha is positively associated with SCZ. Inflammation is believed to play an important role in the pathogenesis of SCZ^44^, and clinical studies have also found rise in inflammatory markers in SCZ^44^. Soluble IL-6 receptor (sIL-6R) may be involved in ‘trans-signaling’, in which the complex of IL-6 and sIL-6R may bind to gp130 and exert pro-inflammatory effects^45^. Another recent MR study also reported increased sIL-6R being associated with SCZ^46^, but here the proteomics dataset used is different and the association is more specific (for the alpha subunit). Of note, an anti-IL-6 monoclonal antibody (Siltuximab) is currently going through phase 1 and 2 clinical trials for the use to treat schizophrenia (NCT02796859), highlighting the translational potential of finding causal proteins for psychiatric disorders.

While the strength of significance was lower, high sIL-6R (involved in trans-signaling of IL-6) was also observed to be causally linked to *higher* MDD (KORA β=0.015, INTERVAL β=0.016, Simes P=0.019) and PTSD risks (KORA β=0.067, INTERVAL β=0.057, Simes P=0.037). Previous studies showed that inflammatory cytokines, such as IL-6, might be positively associated with MDD^73-75^. IL-6 signaling may be related to multiple crucial processes in the pathophysiology of depression^76^. One of the possible pathways is that IL-6, along with other mediators, such as glutamate, neurotrophic factor and infectious agents, activate kappaB kinase (IκK) and controls the downstream NFκB, which is thought to affect synaptic signaling and neural morphology necessary and sufficient for generating depression-like symptoms in chronic social defeat mice models^77^. Of note, NFκB can also induce IL-6 expression^78,79^. In intervention studies, pharmacological intervention on IL-6 level using COX-2 inhibitors reduces depressive symptoms in patients^80-83^. Another pathway by which IL-6 could possibly affect depression is by acting on indoleamine 2,3-dioxygenase (IDO). This increases peripheral KYN, which may then cross the blood brain barrier, contributing to higher central KYN levels^84^. This pathway is further elaborated below in the KYNU→MDD section. Moreover, a metaanalysis by Passo et al. in PTSD patients revealed IL-6 to be one of the consistently elevated cytokines^85^. Interestingly, elevation of IL-6 was still observed after exclusion of subjects with comorbid MDD.

##### The kynurenine pathway

The kynurenine pathway is another pathway which may be of clinical relevance. We found that KYNU positively affect MDD (β=0.131, Simes p=4.16E-3). Notably, KYNU is a target of several drugs including L-tryptophan, melatonin and oxitriptan. Tryptophan is precursor for serotonin synthesis and has long been investigated for treatment of depression and anxiety disorders^86^. There is some evidence for clinical benefit in depression, but the findings are inconclusive. Biologically, KYNU is an enzyme that catalyzes the conversion of 3-hydroxykynurenine (OHK), a metabolite of kynurenine (KYN), to 3-hydroxyanthranilic acid (3-HAA). 3-HAA can be further degraded to quinolinic acid (QUIN) which is considered neurotoxic because of its agonistic action on the NMDA receptor. In an alternative pathway, KYN is converted to kynurenic acid (KYNA). In major depression, elevated QUIN and reduction of KYNA/QUIN ratio has been observed^87^,which was found to be reversed with electroconvulsive therapy^88^. Our findings, coupled with the previous literature, suggest that the kynurenine pathway may play a role in the pathogenesis of depression.

### MR analysis: Psychiatric disorders/traits as exposure and proteins as outcome

#### MR analysis results

We also performed MR in the other direction, with psychiatric disorders/traits as exposure and proteins as outcome. The results are presented in Table S4 (top results in Table 3). In total 16 psychiatric disorders have at least one genetic variant having *p*<5e-8 (Table S4a). However, some of these genome-wide significant SNPs cannot map to the SNP panels of KORA and INTERVAL. Finally, 11 and 15 psychiatric traits remained to be analyzed with KORA and INTERVAL, considering plasma protein levels as outcome.

**Table 3.**
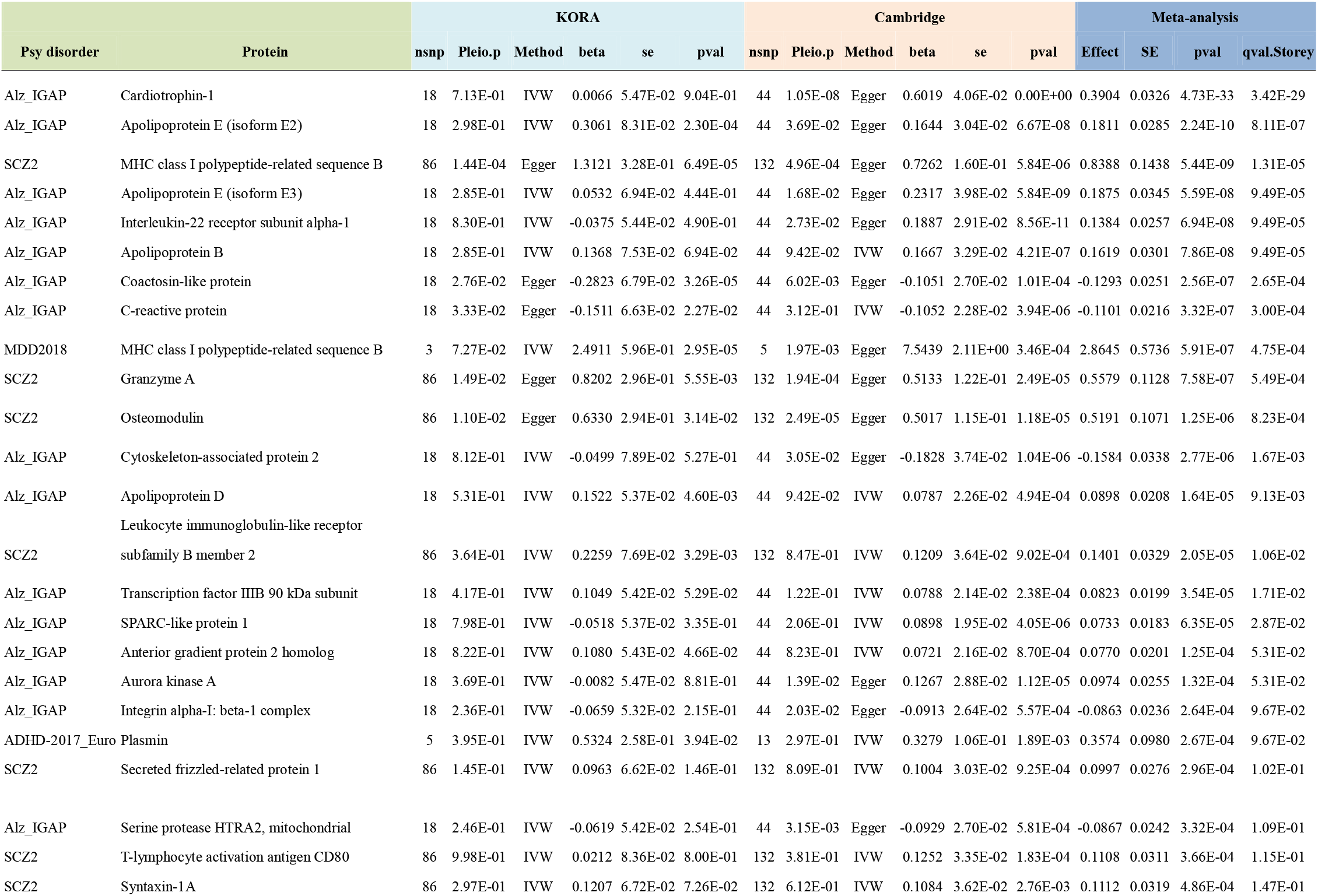
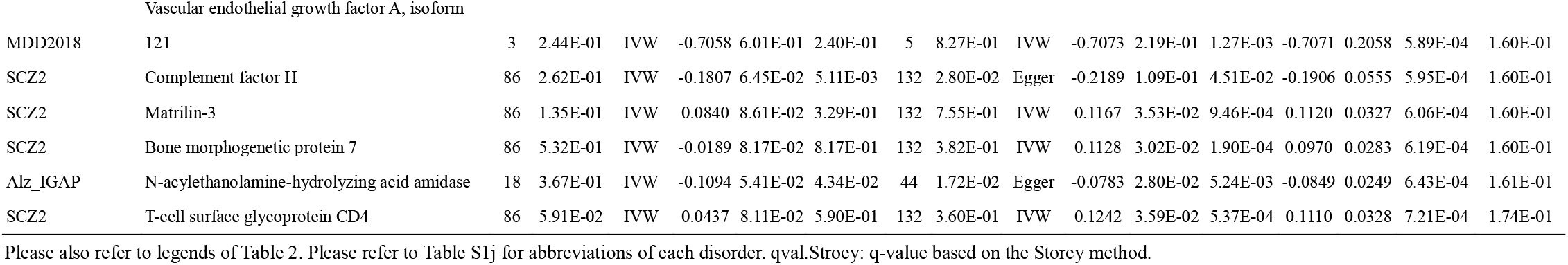
MR results with psychiatric disorders/traits as exposure and proteins as outcome (joint analysis of KORA and INTERVAL)

Totally 12281 and 36108 MR tests were performed based on the KORA and INTERVAL datasets respectively. Note that since KORA and INTERVAL are independent outcome samples, we may consider the analyses as ‘replications’ of one another. Based on the INTERVAL sample alone, 132 disease-protein pairs had FDR<=0.1 and 274 had FDR<=0.2 (Table S4c). The other sample KORA was much smaller in size, and a smaller number of significant findings were observed. In total two disease-protein associations achieved FDR<=0.2 and 10 had FDR<=0.5 (Table S4b). Finally we conducted meta-analysis on the pairs of associations analyzed in both pQTL samples (Table S4d). In total 20, 30 and 145 meta-analysis results achieved FDR <=0.1, <=0.2 and <=0.5 respectively. In occasional cases single-SNP MR has to be performed; HEIDI results are presented alongside the main analysis in Table S4b and S4c.

#### Highlights of top disease-to-protein associations

We observed a number of interesting disease-to-protein associations supported by literature. For the limit of space, we mainly discuss on proteins that are elevated/reduced as a result of SCZ. For example, Secreted Frizzled related protein 1 (sFRP-1) plays a role in WnT signaling, and studies also showed they affect development of dopaminergic neurons in vivo^89^. Wnt-signaling has also implicated in schizophrenia, and some other genes in the pathway, such as DISC1 and GSK3B have been suggested as candidate genes for the disease^90^. Syntaxin-1 plays an important role in synaptic functioning and neurotransmitter release, and disrupted synaptic transmission and connectivity may underlie the pathogenesis of SCZ^91^. We found a potential causal association of SCZ with raised syntaxin-1 A; interestingly, another study also revealed increased syntaxin-1A in SCZ cases compared to controls, which was reduced upon antipsychotic treatment^92^. Another protein BMP7 showed a positive causal relationship with SCZ. Of note, BMP signaling, especially BMP7, were shown in previous studies to regulate dendritic formation in cortical neurons^93, 94^. In another study of differentially expressed genes using post-mortem SCZ brain samples, the authors also found BMP7 as one of the top-ranked genes with higher expression in SCZ subjects^95^. Also, it is worth highlighting that quite a number of the top proteins were related to immune pathways or inflammation. For example, CD80, complement factor H, CD4, interferon lambda-2, Killer cell immunoglobulin-like receptor 2DL5A, leukocyte immunoglobulin-like receptor subfamily B member 2, complement C4 etc. were all ranked among the top (with FDR<0.2) in KORA or INTERVAL samples or meta-analysis. Interestingly, a recent clinical study reported that C4 was significantly increased in chronic SCZ patient and was associated with positive and negative symptom severity {Cropley, 2018 #164}, highlighting the potential of C4 as a disease activity biomarker. Immune system dysfunctioning and neuro-inflammation have long been implicated in SCZ. While immune dysfunction may a cause of schizophrenia^96^, the above findings also suggest that the disease may also trigger inflammation and/or changes in the immune system, which in turn may be related to the prognosis of illness.

### Directed network analysis of protein-disease associations

We constructed a directed network using the bidirectional relationships revealed by MR between proteins and diseases, which are pruned by FDR<0.1 (Figure 2). The directed network provides a clearer visualization on how proteins affecting multiple diseases or one disease without the other. For example, we note that some of the proteins might be associated with more than one disorder, but in different directions. If findings are replicated, such blood proteins may have the potential to serve as biomarkers for differential diagnosis of closely-related conditions. They may also shed light on the difference in pathophysiology of different psychiatric disorders. Taking SCZ and bipolar disorder as example, FTH1 and EPHA1 *decrease* the risk of bipolar disorder (FTH1:β=-0.058, Simes p=0.0611 and EPHA1:β=-0.046, Simes p=0.046) but they *increase* risk of schizophrenia (FTH1:β=0.109, Simes p=0.043; EPHA1:β=0.118, Simes p=0.008). On the other hand, WFIKKN2 increases risk of schizophrenia (β=0.074, Simes p=0.002) while it is protective against bipolar disorder (β=-0.097, Simes p=0.028). Proteins that show causal relationships with solely one disorder but not others may also be useful in distinguishing between disorders. For instance, MICB (β = 1.019, Simes p=1.17e-05), OMD β = 0.567, Simes p = 2.36e-05), GZMA (β=0.667, Simes p=4.98e-05), were elevated solely due to schizophrenia.

**Figure 2.**
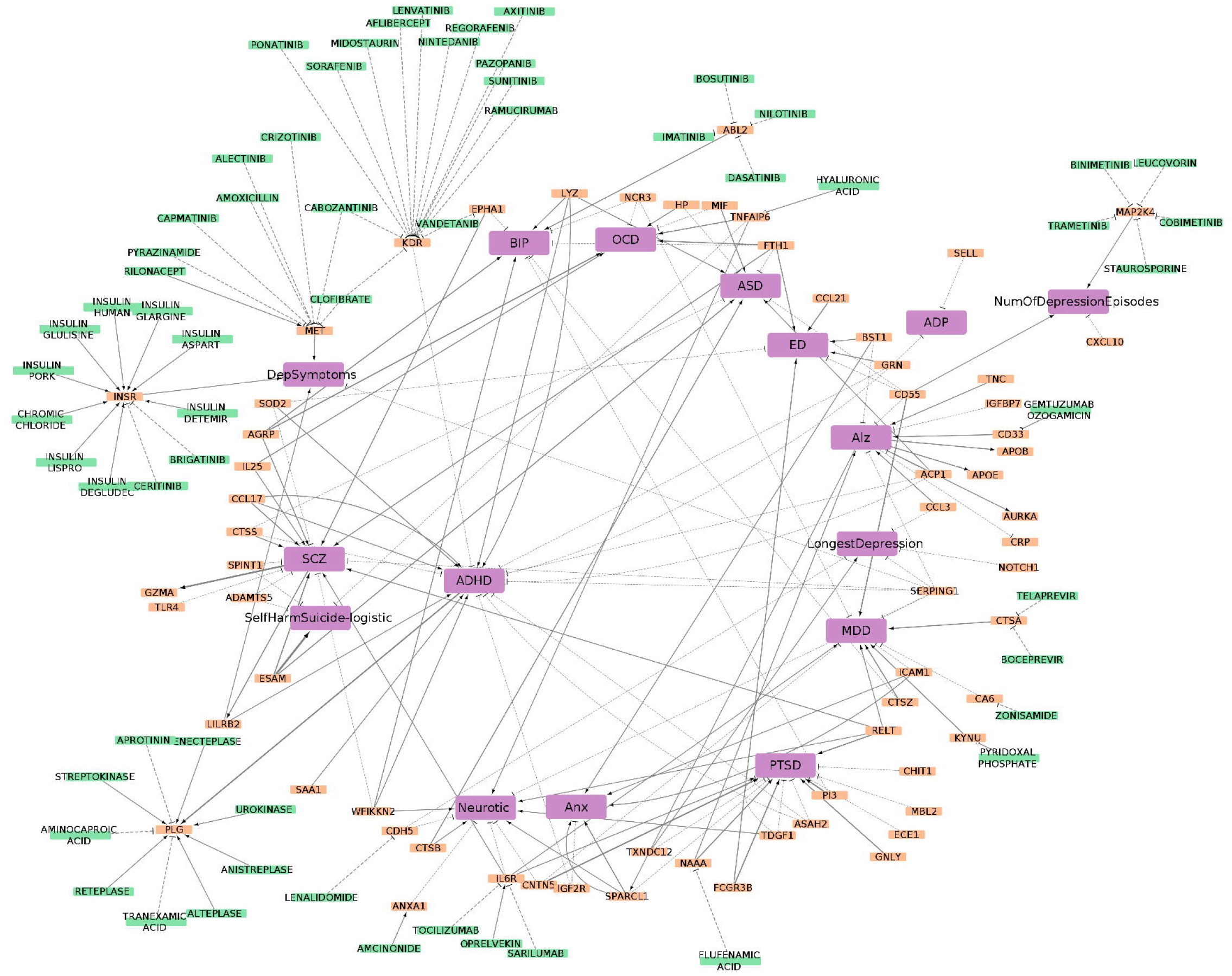
Directed network of proteins, drugs and psychiatric disorders/traits based on MR results (proteinprotein interactions not considered). Here we only considered proteins present in both KORA and INTERVAL samples (with FDR<0.1).

To gain insight and provide further evidence into the relationships between proteins and disease, we performed a systematic literature search to reveal the known relationship between diseases and proteins. Of the 213 significant (FDR < 0.1) causal pairs of protein and disease, 76 were supported by at least one external study. The evidence is visualized with another network with number of evidence as weighting (Figure S1).

### Enriched pathways for the highlighted proteins

We analyzed the functional pathways that are enriched in our directed network to understand the roles of the proteins in causal relationships with psychiatric diseases (Figure S2; Table S5b). Interestingly, many of the top pathways were related to inflammation or immune functioning. The top 5 significant pathways includes: Human Complement System (WP:2806, Bonferroni-corrected pval=1.06E-9), Neutrophil degranulation (R-HSA:6798695, Bonf pval=1.19E-9), Complement cascade (R-HSA: 166658, Bonf pval=4.08E-8), Cytokine-cytokine receptor interaction (KEGG:04060, Bonf pval=7.38E-8), Vitamin B12 Metabolism (WP:1533, Bonf pval=4.60E-7).

## Discussion

Here we have conducted a causal proteome-wide association study of over 3000 plasma proteins with 24 GWAS datasets of psychiatric traits/disorders. We leveraged GWAS summary statistics of plasma protein measurements and psychiatric disorders, and perform bi-directional Mendelian randomization analysis to uncover casual relationships between the two entities. We have performed ~95,000 MR tests in total, and also performed several additional analyses such as HEIDI test, analysis in *cis*-variants only as well as replication in independent datasets (for both protein-to-disease and disease-to-proteins analyses). We believe this work will shed light on the pathophysiology underlying the development and progression of psychiatric disorders. It may also facilitate the development of novel biomarkers for prediction, diagnosis and monitoring of disease progression or recovery. By targeting proteins that may be causal to the disease, our findings may also help drug development or repurposing for psychiatric disorders. Importantly, all our analysis results are also made available online and are easily searchable, which we believe will represent a useful resource for basic and clinical researchers interested in further studies of plasma proteins in psychiatric disorders.

Here we conducted *bi*-directional analyses to investigate not only how proteins affect psychiatric disorder risks, but also how psychiatric traits/disorders may be causal risk factors for changes in protein levels. This is a unique feature of the current study, as previous works on ‘omics’-based MR usually focused on, for example, how expression or methylation changes contribute to diseases^37^. Causal relationships in the reverse direction are however far less explored. By the definition of causality, if an intervention is applied to the cause (disease), then the outcome (protein) should change. Therefore such biomarkers are theoretically justified to evaluate the disease progression, treatment response or resolution from disease symptoms. In this work, we have found a number of putative disease-to-protein associations worthy of further investigations, as already highlighted above.

As a technical note, when the MR exposure is a binary variable (disease or not), it is preferable to conceptualize it as a (possibly latent) continuous risk factor^97^. For example, if the studied exposure is hypertension, then blood pressure would be the underlying continuous risk factor or trait^97^. For most psychiatric disorders, we may also conceptualize a continuous factor underlying the disorder. For instance, depressive symptoms can be the latent factor underlying MDD. Similarly, one can conceptualize a continuous scale of (or tendency to) anxiety, psychosis, OCD etc. as the latent risk factor or trait. The principles of MR causal inference still apply.

In addition to the potential clinical applications described above, we also wish to highlight two other aspects of translational potential. Besides diagnosis-based phenotypes, we also included in our analysis a GWAS on antidepressant response (as both binary and continuous outcome), and a GWAS on SCZ *versus* bipolar disorder (BD) patients. The former analysis aims to reveal proteins causally associated with response to antidepressants, and may help discover candidate protein markers to predict treatment response. MR analysis of the latter dataset (SCZ vs BD) could help to explore differences in proteomic profiles between the two highly related disorders. The results may ultimately help to prioritize protein biomarkers that may contribute to differential diagnosis (DDx) between BD and SCZ, given that response to medications such as lithium and valproate can differ between the disorders. Obviously similar analysis may also be conducted for other kinds of psychiatric disorders, hence the presented analytic framework also points to a new direction for finding biomarkers in predicting drug response and DDx in general.

Another contribution of this study is a directed network analysis of disease protein associations of psychiatric traits/disorders. A network approach to analyzing genetic and protein associations enables us to gain more comprehensive insight into the pathways and complex pathophysiology underlying complex diseases. Several methods have been developed for network analysis of GWAS data^98, 99^. However, previous analytic methods have mainly focused on *un*directed networks; to our knowledge, this is the first study to present a *directed* network analysis framework across *multiple* complex disease traits, based on GWAS data and the principle of MR. Such networks are useful in unraveling the mechanisms underlying disease comorbidities and complications.

There are several limitations that are worth noting. The MR approach is less vulnerable to reverse causality and confounding compared to observational studies, however in some scenarios it is still not easy to firmly establish causality. In the study of causal effects of proteins to psychiatric disorders, the number of instrument SNPs is usually quite small, and it is quite common to rely on only one instrument (i.e. single-SNP MR). There are no methods to distinguish between causality and pleiotropy in single-SNP MR, although the HEIDI test can help to identify ‘linkage’ from the same SNP being associated with protein levels and disease at the same time^37^. When the number of instruments is larger, more tools are available to address the assumptions of MR, especially pleiotropy^57^. However, the number of instruments is still typically small when proteins are treated as exposure; the power of methods like Egger regression to detect pleiotropy is relatively weak under such scenarios^33^. Methods based on outlier removal (e.g. MR-PRESSO^100^) are also not applicable in the presence of few instruments. On the whole, we could still prioritize proteins that are more likely causal to diseases, although fully establishing causality remains difficult. Nevertheless, at least the current study could show which proteins may be *associated with* various psychiatric disorders, which helps to limit the search space for suitable protein biomarkers or target proteins for drug development. For MR analysis with psychiatric disorders as exposure, the above problem of inadequate instruments is usually of a lesser concern, especially for disorders like schizophrenia and Alzheimer’s disease with large GWAS sample sizes and multiple genetic instruments.

The MR approach itself also has limitations. As MR is based on genetic instruments to represent the exposure, the results reflect a chronic exposure of (genetically predicted) higher/lower protein levels to the psychiatric disorder. The effects of short-term exposure, such as taking an inhibitor targeting the protein, may not be the same as effects from long-term exposure as predicted by MR.

With regards to the study samples, many psychiatric disorders are highly heterogeneous with regards to their presentations, course of disease, prognosis etc^101^. A protein may be associated with a subgroup of patients with certain clinical characteristics only (e.g. female patients with early onset) but not with the other groups. A future direction is to conduct the analysis in stratified or more deeply-phenotyped samples (e.g. male or female-only; early or late-onset only; with a particular clinical presentation). Also, for some of the samples (especially those from UKBB), the disease diagnosis or symptoms are self-reported instead of being assessed by a clinical professional. The latter is probably not practical in a biobank due to high cost of performing detailed assessments. Nevertheless, further replications in more carefully assessed and phenotyped samples are warranted.

Also related to sample characteristics, we have tried to employ the latest and largest GWAS samples. Nevertheless, for some of the psychiatric disorders (e.g. OCD or PTSD), the sample size may still be relatively small. Also we have performed replication analysis in the UKBB, however the UKBB in general does not contain a lot of patients with psychiatric disorders (except for depression and anxiety). As a result, the power of replication study is relatively limited and absence of significant replications does not exclude genuine associations.

One potential translational aspect is the development of blood-based biomarkers for psychiatric disorders. While this work we mainly study the protein candidates one by one, in practice and in future studies, we may combine different biomarkers and even other modalities (e.g. neuroimaging) to improve the performance of the test. This will be left as a direction for future investigations.

In summary, we have conducted a proteome-wide association study to decipher the causal relationship between a large panel of plasma proteins and psychiatric disorders. We believe this work sheds light on the pathophysiology of psychiatric disorders, and will help facilitate the development of protein biomarkers as well as new or repurposed drugs for the treatment of psychiatric disorders.

## Supporting information

Supplementary Note on all tables

## Acknowledgements

We would like to thank Prof. Stephen Tsui and the Hong Kong Bioinformatics Center for computing support. This study was partially supported by the Lo Kwee Seong Biomedical Research Fund, an NSFC grant (81971706), the Health and Medical Research Fund and a Chinese University of Hong Kong Direct Grant.

## Author contributions

Conceived and designed the study: HCS. Supervised the study: HCS. Main data analysis: CKLC, ALCL.

Data interpretation: HCS, CKLC, ALCL, PCS. Data analysis methodology: HCS (lead), ALCL with input from PCS. Protein Network analysis: ALCL. Drafted the manuscript: HCS (lead), with input from ALCL.

## Figure Legends

Figure S1 Undirected network based on MR relationship literature evidence available on PubMed. Thicker edges indicate more matches in PubMed between the disease-protein pair. Direction of effect was not considered.

Figure S2 Pathway enrichment of the proteins in significant (FDR<0.1) disease-protein pairs. This functional annotation is performed with ClueGO using KEGG (Release 89.1), Reactome Pathways (ver. 67) and WikiPathways (February 2019 Release) as reference background datasets. Significance (as indicated by bubble size) of enrichment is estimated with hypergeometric tests.

## References

1. Vigo D, Thornicroft G, Atun R. Estimating the true global burden of mental illness. Lancet Psychiatry 2016; 3(2): 171–178.

2. Venkatasubramanian G, Keshavan MS. Biomarkers in Psychiatry - A Critique. Ann Neurosci 2016; 23(1): 3–5.

3. Mora C, Zonca V, Riva MA, Cattaneo A. Blood biomarkers and treatment response in major depression. Expert Rev Mol Diagn 2018; 18(6): 513–529.

4. Comes AL, Papiol S, Mueller T, Geyer PE, Mann M, Schulze TG. Proteomics for blood biomarker exploration of severe mental illness: pitfalls of the past and potential for the future. Transl Psychiatry 2018; 8(1): 160.

5. Bennett DA, Holmes MV. Mendelian randomisation in cardiovascular research: an introduction for clinicians. Heart 2017.

6. Davey Smith G, Hemani G. Mendelian randomization: genetic anchors for causal inference in epidemiological studies. Hum Mol Genet 2014; 23(R1): R89–98.

7. Johnson EC, Border R, Melroy-Greif WE, de Leeuw CA, Ehringer MA, Keller MC. No Evidence That Schizophrenia Candidate Genes Are More Associated With Schizophrenia Than Noncandidate Genes. Biol Psychiatry 2017; 82(10): 702–708.

8. Border R, Johnson EC, Evans LM, Smolen A, Berley N, Sullivan PF et al. No Support for Historical Candidate Gene or Candidate Gene-by-Interaction Hypotheses for Major Depression Across Multiple Large Samples. Am J Psychiatry 2019; 176(5): 376–387.

9. Zhang F, Lupski JR. Non-coding genetic variants in human disease. Human Molecular Genetics 2015; 24: R102–R110.

10. Scott KM. Depression, anxiety and incident cardiometabolic diseases. Curr Opin Psychiatry 2014; 27(4): 289–293.

11. Wong BC, Chau CK, Ao FK, Mo CH, Wong SY, Wong YH et al. Differential associations of depression-related phenotypes with cardiometabolic risks: Polygenic analyses and exploring shared genetic variants and pathways. Depress Anxiety 2019; 36(4): 330–344.

12. Halaris A. Inflammation-Associated Co-morbidity Between Depression and Cardiovascular Disease. Curr Top Behav Neurosci 2017; 31: 45–70.

13. Suhre K, Arnold M, Bhagwat AM, Cotton RJ, Engelke R, Raffler J et al. Connecting genetic risk to disease end points through the human blood plasma proteome. Nat Commun 2017; 8: 14357.

14. Sun BB, Maranville JC, Peters JE, Stacey D, Staley JR, Blackshaw J et al. Genomic atlas of the human plasma proteome. Nature 2018; 558(7708): 73–79.

15. Martin J, Walters RK, Demontis D, Mattheisen M, Lee SH, Robinson E et al. A Genetic Investigation of Sex Bias in the Prevalence of Attention-Deficit/Hyperactivity Disorder. Biol Psychiatry 2018; 83(12): 1044-1053.

16. Duncan L, Yilmaz Z, Gaspar H, Walters R, Goldstein J, Anttila V et al. Significant Locus and Metabolic Genetic Correlations Revealed in Genome-Wide Association Study of Anorexia Nervosa. Am J Psychiatry 2017; 174(9): 850–858.

17. Lambert JC, Ibrahim-Verbaas CA, Harold D, Naj AC, Sims R, Bellenguez C et al. Meta-analysis of 74,046 individuals identifies 11 new susceptibility loci for Alzheimer’s disease. Nat Genet 2013; 45(12): 1452-1458.

18. Grove J, Ripke S, Als TD, Mattheisen M, Walters RK, Won H et al. Identification of common genetic risk variants for autism spectrum disorder. Nat Genet 2019; 51(3): 431–444.

19. Otowa T, Hek K, Lee M, Byrne EM, Mirza SS, Nivard MG et al. Meta-analysis of genome-wide association studies of anxiety disorders. Mol Psychiatr 2016; 21(10): 1391–1399.

20. Okbay A, Baselmans BM, De Neve JE, Turley P, Nivard MG, Fontana MA et al. Genetic variants associated with subjective well-being, depressive symptoms, and neuroticism identified through genomewide analyses. Nat Genet 2016; 48(6): 624–633.

21. Wray NR, Ripke S, Mattheisen M, Trzaskowski M, Byrne EM, Abdellaoui A et al. Genome-wide association analyses identify 44 risk variants and refine the genetic architecture of major depression. Nat Genet 2018; 50(5): 668–681.

22. consortium C. Sparse whole-genome sequencing identifies two loci for major depressive disorder. Nature 2015; 523(7562): 588–591.

23. UK Biobank results from Neale’s Lab.

24. International Obsessive Compulsive Disorder Foundation Genetics C, Studies OCDCGA. Revealing the complex genetic architecture of obsessive-compulsive disorder using meta-analysis. Mol Psychiatry 2018; 23(5): 1181–1188.

25. Duncan LE, Ratanatharathorn A, Aiello AE, Almli LM, Amstadter AB, Ashley-Koch AE et al. Largest GWAS of PTSD (N=20 070) yields genetic overlap with schizophrenia and sex differences in heritability. Mol Psychiatry 2018; 23(3): 666–673.

26. Stahl EA, Breen G, Forstner AJ, McQuillin A, Ripke S, Trubetskoy V et al. Genome-wide association study identifies 30 loci associated with bipolar disorder. Nature Genetics 2019; 51(5): 793–+.

27. Schizophrenia Working Group of the Psychiatric Genomics C. Biological insights from 108 schizophrenia-associated genetic loci. Nature 2014; 511(7510): 421–427.

28. Investigators G, Investigators M, Investigators SD. Common genetic variation and antidepressant efficacy in major depressive disorder: a meta-analysis of three genome-wide pharmacogenetic studies. Am J Psychiatry 2013; 170(2): 207–217.

29. Ruderfer DM, Fanous AH, Ripke S, McQuillin A, Amdur RL, Schizophrenia Working Group of the Psychiatric Genomics C et al. Polygenic dissection of diagnosis and clinical dimensions of bipolar disorder and schizophrenia. Mol Psychiatry 2014; 19(9): 1017–1024.

30. Ramtekkar UP, Reiersen AM, Todorov AA, Todd RD. Sex and age differences in attention-deficit/hyperactivity disorder symptoms and diagnoses: implications for DSM-V and ICD-11. J Am Acad Child Adolesc Psychiatry 2010; 49(3): 217–228 e211-213.

31. Hourani L, Williams J, Bray R, Kandel D. Gender differences in the expression of PTSD symptoms among active duty military personnel. J Anxiety Disord 2015; 29:101–108.

32. Burgess S, Dudbridge F, Thompson SG. Combining information on multiple instrumental variables in Mendelian randomization: comparison of allele score and summarized data methods. Stat Med 2016; 35(11): 1880–1906.

33. Bowden J, Smith GD, Burgess S. Mendelian randomization with invalid instruments: effect estimation and bias detection through Egger regression. Int J Epidemiol 2015; 44(2): 512–525.

34. Benjamini Y, Hochberg Y. Controlling the False Discovery Rate - a Practical and Powerful Approach to Multiple Testing. J Roy Stat Soc B Met 1995; 57(1): 289–300.

35. Storey JD. The positive false discovery rate: A Bayesian interpretation and the q-value. Ann Stat 2003; 31(6): 2013–2035.

36. Benjamini Y, Yekutieli D. The control of the false discovery rate in multiple testing under dependency. Ann Stat 2001; 29(4): 1165–1188.

37. Zhu Z, Zhang F, Hu H, Bakshi A, Robinson MR, Powell JE et al. Integration of summary data from GWAS and eQTL studies predicts complex trait gene targets. Nat Genet 2016; 48(5): 481–487.

38. Nichols T, Brett M, Andersson J, Wager T, Poline JB. Valid conjunction inference with the minimum statistic. Neuroimage 2005; 25(3): 653–660.

39. Simes RJ. An Improved Bonferroni Procedure for Multiple Tests of Significance. Biometrika 1986; 73(3): 751–754.

40. Heller R, Bogomolov M, Benjamini Y. Deciding whether follow-up studies have replicated findings in a preliminary large-scale omics study. P Natl Acad Sci USA 2014; 111(46): 16262–16267.

41. Lloyd-Jones LR, Robinson MR, Yang J, Visscher PM. Transformation of Summary Statistics from Linear Mixed Model Association on All-or-None Traits to Odds Ratio. Genetics 2018; 208(4): 1397–1408.

42. Cotto KC, Wagner AH, Feng YY, Kiwala S, Coffman AC, Spies G et al. DGIdb 3.0: a redesign and expansion of the drug-gene interaction database. Nucleic acids research 2018; 46(D1): D1068–d1073.

43. Wang H, Yin YX, Gong DM, Hong LJ, Wu G, Jiang Q et al. Cathepsin B inhibition ameliorates leukocyte-endothelial adhesion in the BTBR mouse model of autism. Cns Neurosci Ther 2019; 25(4): 476–485.

44. Khandaker GM, Cousins L, Deakin J, Lennox BR, Yolken R, Jones PB. Inflammation and immunity in schizophrenia: implications for pathophysiology and treatment. Lancet Psychiatry 2015; 2(3): 258–270.

45. Rose-John S. IL-6 Trans-Signaling via the Soluble IL-6 Receptor: Importance for the Pro-Inflammatory Activities of IL-6. Int J Biol Sci 2012; 8(9): 1237–1247.

46. Hartwig FP, Borges MC, Horta BL, Bowden J, Smith GD. Inflammatory Biomarkers and Risk of Schizophrenia A 2-Sample Mendelian Randomization Study. Jama Psychiat 2017; 74(12): 1226–1233.

47. Kern JK, Geier DA, Sykes LK, Geier MR. Relevance of Neuroinflammation and Encephalitis in Autism. Front Cell Neurosci 2016; 9.

48. Kealy J, Greene C, Campbell M. Blood-brain barrier regulation in psychiatric disorders. Neurosci Lett 2018:133664.

49. Najjar S, Pahlajani S, De Sanctis V, Stern JNH, Najjar A, Chong D. Neurovascular Unit Dysfunction and Blood-Brain Barrier Hyperpermeability Contribute to Schizophrenia Neurobiology: A Theoretical integration of Clinical and experimental evidence. Front Psychiatry 2017; 8.

50. Yamauchi N, Shibasaki T, Wakabayashi I, Demura H. Brain beta-endorphin and other opioids are involved in restraint stress-induced stimulation of the hypothalamic-pituitary-adrenal axis, the sympathetic nervous system, and the adrenal medulla in the rat. Brain Res 1997; 777(1-2): 140–146.

51. Iyengar S, Kim HS, Wood PL. Mu-Opioid, Delta-Opioid, Kappa-Opioid and Epsilon-Opioid Receptor Modulation of the Hypothalamic Pituitary Adrenocortical (Hpa) Axis - Subchronic Tolerance Studies of Endogenous Opioid-Peptides. Brain Res 1987; 435(1-2): 220–226.

52. Hegadoren KM, O’Donnell T, Lanius R, Coupland NJ, Lacaze-Masmonteil N. The role of beta-endorphin in the pathophysiology of major depression. Neuropeptides 2009; 43(5): 341–353.

53. Latif A, Heinz P, Cook R. Iron deficiency in autism and Asperger syndrome. Autism 2002; 6(1): 103–114.

54. Herguner S, Kelesoglu FM, Tanidir C, Copur M. Ferritin and iron levels in children with autistic disorder. Eur J Pediatr 2012; 171(1): 143–146.

55. Dosman CF, Brian JA, Drmic IE, Senthilselvan A, Harford MM, Smith RW et al. Children with autism: Effect of iron supplementation on sleep and ferritin. Pediatr Neurol 2007; 36(3): 152–158.

56. Marioni RE, Harris SE, Zhang Q, McRae AF, Hagenaars SP, Hill WD et al. GWAS on family history of Alzheimer’s disease. Transl Psychiatry 2018; 8(1): 99.

57. Hemani G, Bowden J, Smith GD. Evaluating the potential role of pleiotropy in Mendelian randomization studies. Human Molecular Genetics 2018; 27(R2): R195–R208.

58. Jansen PM, Eisele B, de Jong IW, Chang A, Delvos U, Taylor FB, Jr. et al. Effect of C1 inhibitor on inflammatory and physiologic response patterns in primates suffering from lethal septic shock. Journal of immunology 1998; 160(1): 475–484.

59. Bradshaw EM, Chibnik LB, Keenan BT, Ottoboni L, Raj T, Tang A et al. CD33 Alzheimer’s disease locus: altered monocyte function and amyloid biology. Nat Neurosci 2013; 16(7): 848–U892.

60. Jiang T, Yu JT, Hu N, Tan MS, Zhu XC, Tan L. CD33 in Alzheimer’s Disease. Mol Neurobiol 2014; 49(1): 529–535.

61. Lowin T, Apitz M, Anders S, Straub RH. Anti-inflammatory effects of N-acylethanolamines in rheumatoid arthritis synovial cells are mediated by TRPV1 and TRPA1 in a COX-2 dependent manner. Arthritis Res Ther 2015; 17: 321–321.

62. Pistis M, Muntoni AL. Roles of N-Acylethanolamines in Brain Functions and Neuropsychiatric Diseases. In: Melis M (ed). Endocannabinoids and Lipid Mediators in Brain Functions. Springer International Publishing: Cham, 2017, pp 319–346.

63. Ueda N, Tsuboi K, Uyama T. Metabolism of endocannabinoids and related N-acylethanolamines: canonical and alternative pathways. The FEBS journal 2013; 280(9): 1874–1894.

64. Sun YX, Tsuboi K, Zhao LY, Okamoto Y, Lambert DM, Ueda N. Involvement of N-acylethanolamine-hydrolyzing acid amidase in the degradation of anandamide and other N-acylethanolamines in macrophages. Biochimica et biophysica acta 2005; 1736(3): 211–220.

65. Koppel J, Davies P. Targeting the endocannabinoid system in Alzheimer’s disease. J Alzheimers Dis 2008; 15(3): 495–504.

66. Bitencourt RM, Takahashi RN. Cannabidiol as a Therapeutic Alternative for Post-traumatic Stress Disorder: From Bench Research to Confirmation in Human Trials. Frontiers in neuroscience 2018; 12: 502.

67. Berardi A, Schelling G, Campolongo P. The endocannabinoid system and Post Traumatic Stress Disorder (PTSD): From preclinical findings to innovative therapeutic approaches in clinical settings. Pharmacological research 2016; 111: 668–678.

68. Hill MN, Campolongo P, Yehuda R, Patel S. Integrating Endocannabinoid Signaling and Cannabinoids into the Biology and Treatment of Posttraumatic Stress Disorder. Neuropsychopharmacology: official publication of the American College of Neuropsychopharmacology 2018; 43(1): 80–102.

69. Pinna G. Biomarkers for PTSD at the Interface of the Endocannabinoid and Neurosteroid Axis. Frontiers in neuroscience 2018; 12: 482–482.

70. Fraser GA. The use of a synthetic cannabinoid in the management of treatment-resistant nightmares in posttraumatic stress disorder (PTSD). Cns Neurosci Ther 2009; 15(1): 84–88.

71. Bottemanne P, Muccioli GG, Alhouayek M. N-acylethanolamine hydrolyzing acid amidase inhibition: tools and potential therapeutic opportunities. Drug Discovery Today 2018; 23(8): 1520–1529.

72. Beggiato S, Tomasini MC, Ferraro L. Palmitoylethanolamide (PEA) as a Potential Therapeutic Agent in Alzheimer’s Disease. Front Pharmacol 2019; 10: 821–821.

73. Valkanova V, Ebmeier KP, Allan CL. CRP, IL-6 and depression: a systematic review and meta-analysis of longitudinal studies. J Affect Disord 2013; 150(3): 736–744.

74. Hodes GE, Ménard C, Russo SJ. Integrating Interleukin-6 into depression diagnosis and treatment. Neurobiol Stress 2016; 4: 15–22.

75. Himmerich H, Patsalos O, Lichtblau N, Ibrahim MAA, Dalton B. Cytokine Research in Depression: Principles, Challenges, and Open Questions. Front Psychiatry 2019; 10: 30.

76. Anderson G, Kubera M, Duda W, Lason W, Berk M, Maes M. Increased IL-6 trans-signaling in depression: focus on the tryptophan catabolite pathway, melatonin and neuroprogression. Pharmacological reports: PR 2013; 65(6): 1647–1654.

77. Christoffel DJ, Golden SA, Heshmati M, Graham A, Birnbaum S, Neve RL et al. Effects of inhibitor of kappaB kinase activity in the nucleus accumbens on emotional behavior. Neuropsychopharmacology: official publication of the American College of Neuropsychopharmacology 2012; 37(12): 2615–2623.

78. Brasier AR. The nuclear factor-kappaB-interleukin-6 signalling pathway mediating vascular inflammation. Cardiovascular research 2010; 86(2): 211–218.

79. Liu T, Zhang L, Joo D, Sun SC. NF-kappaB signaling in inflammation. Signal transduction and targeted therapy 2017; 2.

80. Adzic M, Brkic Z, Mitic M, Francija E, Jovicic MJ, Radulovic J et al. Therapeutic Strategies for Treatment of Inflammation-related Depression. Current neuropharmacology 2018; 16(2): 176–209.

81. Abbasi SH, Hosseini F, Modabbernia A, Ashrafi M, Akhondzadeh S. Effect of celecoxib add-on treatment on symptoms and serum IL-6 concentrations in patients with major depressive disorder: randomized double-blind placebo-controlled study. J Affect Disord 2012; 141(2-3): 308–314.

82. Muller N, Schwarz MJ, Dehning S, Douhe A, Cerovecki A, Goldstein-Muller B et al. The cyclooxygenase-2 inhibitor celecoxib has therapeutic effects in major depression: results of a double-blind,randomized, placebo controlled, add-on pilot study to reboxetine. Mol Psychiatry 2006; 11(7): 680–684.

83. Müller N. COX-2 Inhibitors, Aspirin, and Other Potential Anti-Inflammatory Treatments for Psychiatric Disorders. Front Psychiatry 2019; 10: 375–375.

84. Allison DJ, Ditor DS. The common inflammatory etiology of depression and cognitive impairment: a therapeutic target. J Neuroinflammation 2014; 11: 151.

85. Passos IC, Vasconcelos-Moreno MP, Costa LG, Kunz M, Brietzke E, Quevedo J et al. Inflammatory markers in post-traumatic stress disorder: a systematic review, meta-analysis, and meta-regression. Lancet Psychiatry 2015; 2(11): 1002–1012.

86. Shaw K, Turner J, Del Mar C. Are tryptophan and 5-hydroxytryptophan effective treatments for depression? A meta-analysis. The Australian and New Zealand journal of psychiatry 2002; 36(4): 488–491.

87. Reus GZ, Jansen K, Titus S, Carvalho AF, Gabbay V, Quevedo J. Kynurenine pathway dysfunction in the pathophysiology and treatment of depression: Evidences from animal and human studies. Journal of Psychiatric Research 2015; 68: 316–328.

88. Schwieler L, Samuelsson M, Frye MA, Bhat M, Schuppe-Koistinen I, Jungholm O et al. Electroconvulsive therapy suppresses the neurotoxic branch of the kynurenine pathway in treatment-resistant depressed patients. J Neuroinflamm 2016; 13.

89. Kele J, Andersson ER, Villaescusa JC, Cajanek L, Parish CL, Bonilla S et al. SFRP1 and SFRP2 dose-dependently regulate midbrain dopamine neuron development in vivo and in embryonic stem cells. Stem cells (Dayton, Ohio) 2012; 30(5): 865–875.

90. Inestrosa NC, Montecinos-Oliva C, Fuenzalida M. Wnt Signaling: Role in Alzheimer Disease and Schizophrenia. Journal of Neuroimmune Pharmacology 2012; 7(4): 788–807.

91. Johnson RD, Oliver PL, Davies KE. SNARE proteins and schizophrenia: linking synaptic and neurodevelopmental hypotheses. Acta biochimica Polonica 2008; 55(4): 619–628.

92. Gil-Pisa I, Munarriz-Cuezva E, Ramos-Miguel A, Uriguen L, Meana JJ, Garcia-Sevilla JA. Regulation of munc18-1 and syntaxin-1A interactive partners in schizophrenia prefrontal cortex: down-regulation of munc18-1a isoform and 75 kDa SNARE complex after antipsychotic treatment. The international journal of neuropsychopharmacology 2012; 15(5): 573–588.

93. Horbinski C, Stachowiak EK, Chandrasekaran V, Miuzukoshi E, Higgins D, Stachowiak MK. Bone morphogenetic protein-7 stimulates initial dendritic growth in sympathetic neurons through an intracellular fibroblast growth factor signaling pathway. Journal of neurochemistry 2002; 80(1): 54–63.

94. Guo X, Rueger D, Higgins D. Osteogenic protein-1 and related bone morphogenetic proteins regulate dendritic growth and the expression of microtubule-associated protein-2 in rat sympathetic neurons. Neurosci Lett 1998; 245(3): 131–134.

95. Pietersen CY, Mauney SA, Kim SS, Lim MP, Rooney RJ, Goldstein JM et al. Molecular profiles of pyramidal neurons in the superior temporal cortex in schizophrenia. J Neurogenet 2014; 28(1-2): 53–69.

96. Müller N, Schwarz MJ. Immune System and Schizophrenia. Curr Immunol Rev 2010; 6(3): 213–220.

97. Burgess S, Labrecque JA. Mendelian randomization with a binary exposure variable: interpretation and presentation of causal estimates. Eur J Epidemiol 2018; 33(10): 947–952.

98. Leiserson MD, Eldridge JV, Ramachandran S, Raphael BJ. Network analysis of GWAS data. Current opinion in genetics & development 2013; 23(6): 602–610.

99. Jia P, Zhao Z. Network.assisted analysis to prioritize GWAS results: principles, methods and perspectives. Human genetics 2014; 133(2): 125–138.

100. Verbanck M, Chen CY, Neale B, Do R. Detection of widespread horizontal pleiotropy in causal relationships inferred from Mendelian randomization between complex traits and diseases. Nature Genetics 2018; 50(5): 693–+.

101. Wardenaar KJ, de Jonge P. Diagnostic heterogeneity in psychiatry: towards an empirical solution. Bmc Med 2013; 11.

