## Supplementary Note on all tables for "Uncovering bi-directional causal relationships between plasma proteins and psychiatric disorders: A proteome-wide study and directed network analysis"

This note describes all the supplementary tables. **There are totally 23 supplementary tables, organized into 5 big tables**. Each big table contains a number of tabs ordered by letters.

**Table S1 (7 sub-tables)**

As a whole, Table S1 summarizes the MR results when plasma proteins were treated as exposure and psychiatric disorders/traits as outcome.

S1a – This table summarizes all the pQTLs from the KORA sample.

S1b – This table summarizes all the pQTLs from the INTERVAL sample.

S1c – This table includes all two-sample MR results based on the KORA sample, with plasma protein as exposure and psychiatric disorder/trait as outcome. Please refer to the R package ‘MendelianRandomization’ (<https://cran.r-project.org/web/packages/MendelianRandomization/MendelianRandomization.pdf>) for the output headings. For easy comparison, we also show results of *cis*-only analysis side-by-side.

In addition, this table shows HEIDI results for proteins with FDR (i.e. “q-value”) <0.2 based on Storey’s method. Results are sorted based on *p-*values from the MR analysis.

Heter.pval, heterogeneity p-value; qval, q-value; RSE, estimated residual standard error from the regression model; qval.BH, q-value based on the Benjamini-Hochberg approach; qval.Storey, q-value based on the Storey’s method.

S1d – This table includes all two-sample MR results based on the INTERVAL sample, with plasma protein as exposure and psychiatric disorder/trait as outcome. Results of *cis*-only analysis and HEIDI are shown side-by-side. Results are sorted based on *p*-values from the MR analysis.

S1e – Joint analysis of the KORA and INTERVAL samples, with plasma protein as exposure and psychiatric disorder/trait as outcome. HEIDI results based on the KORA or the INTERVAL sample are included side-by-side for easy reference. HEIDI p>0.05 was considered as a pass. Joint analysis p-values Simes_p and max_p are also presented (see the main text for details). We used the R package ‘metagen’ for the IVW meta-analysis. Please refer to the package documentation (<https://www.rdocumentation.org/packages/meta/versions/4.9-2/topics/metagen>) for the output headings.

Note that results are sorted based on Simes *p*-value of the joint analysis.

S1f – This table shows drugs targeted at selected top proteins showing significant (causal) associations with psychiatric traits in the joint analysis (q-value by Storey method <~0.1 and HEIDI test passed). The list is based on a search against the ChEMBL database.

S1g – Summary of the sources of psychiatric GWAS data. (References are given in the main text)

**Table S2 (5 sub-tables)**

Table S2 summarizes the MR results when plasma proteins were treated as exposure and psychiatric disorders/traits as outcome, but the analysis was performed with ***cis-*pQTLs only.**

S2a – List of cis-eQTLs from the KORA sample

S2b – List of cis-eQTLS from the INTERVAL sample

S2c - MR results based on the KORA sample using ***cis-*pQTLs only**, with plasma protein as exposure and psychiatric disorder/trait as outcome. The table also shows HEIDI results for proteins with FDR (i.e. “q-value”) <0.2 based on Storey’s method.

Results are sorted based on *p-*values from the MR analysis.

S2d –MR results based on the INTERVAL sample using ***cis-*pQTLs only**, with plasma protein as exposure and psychiatric disorder/trait as outcome. The table also shows HEIDI results for proteins with FDR (i.e. “q-value”) <0.2 based on Storey’s method.

Results are sorted based on *p-*values from the MR analysis.

S2e –Joint analysis of the KORA and INTERVAL samples, with plasma protein as exposure and psychiatric disorder/trait as outcome using ***cis-*pQTLs only.** Results are sorted based on Simes *p*-value of the joint analysis. Please also refer to the descriptions of Table S1e.

**Table S3 (5 sub-tables)**

Table S3 summarizes replication analysis of MR results in the UK Biobank (UKBB).

S3a – This table shows how the psychiatric traits from the primary analysis (mainly from the PGC download site, for details see Table S1j) can be ‘mapped’ to UKBB phenotypes. We allow the same primary psychiatric trait to be mapped to multiple UKBB phenotypes.

S3b – This table shows replication of KORA MR results in the UKBB (restricted to protein-disease pairs with FDR<0.1 in the primary analysis based on KORA). Replicability FDR is also presented based on Heller et al. (PNAS, 111(46), 16262-16267). Results are sorted by one-tailed p-value in the UKBB replication sample.

S3c – This table shows replication of INTERVAL MR results in the UKBB (restricted to protein-disease pairs with FDR<0.1 in the primary analysis based on INTERVAL). Results are sorted by one-tailed p-value in the UKBB replication sample.

S3d – Replication in UKBB for a set of **overlapping** proteins between the 2 pQTL datasets with estimated FDR<0.1 in the *joint* analysis by Simes *p*. This table shows analysis based on *KORA pQTL data only* (i.e. INTERVAL dataset not used).

S3e – Replication in UKBB for a set of **overlapping** proteins between the 2 pQTL datasets with estimated FDR<0.1 in the *joint* analysis by Simes *p*. This table shows analysis based on *INTERVAL pQTL data only* (i.e. KORA dataset not used).

**Table S4 (4 sub-tables)**

As a whole, Table S4 summarizes the MR results when psychiatric disorders/traits were treated as exposure and plasma proteins as outcome.

S4a -All two-sample MR results, with psychiatric disorders/traits as exposure and plasma proteins from KORA as outcome. HEIDI results (performed for single-SNP MR with FDR<0.2) are also shown if available. The results are sorted by MR p-values.

S4b - All two-sample MR results, with psychiatric disorders/traits as exposure and plasma proteins from the INTERVAL sample as outcome. HEIDI results (performed for single-SNP MR with FDR<0.2) are also shown if available. The results are sorted by MR p-values.

S4c – Meta-analysis of MR results (psychiatric disorder as exposure and protein as outcome) across the two outcome datasets, KORA and INTERVAL. The results are sorted by the IVW meta-analysis p-values.

S4d - Summary of the number of SNPs passing genome-wide significance in each psychiatric GWAS dataset, and number of SNPs that can be mapped to the genotyping panel of KORA and INTERVAL (after clumping at r^2^ = 0.1)

**Table S5 (2 sub-tables)**

Table S5a	Relevant drug matches in DGIdb for top proteins from MR analysis

Table S5b	Gene ontology (GO) enrichment analysis results of top proteins identified in MR (that are also shown in the network)
