## Supplementary Text for "Uncovering bi-directional causal relationships between plasma proteins and psychiatric disorders: A proteome-wide study and directed network analysis"

*MR analysis methods*

The IVW framework is very widely used in MR studies. Here we used an IVW approach that is able to account for SNP correlations as described in Burgess et al.^1^. Briefly, assume
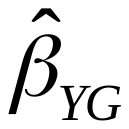
 to be the vector of estimated regression coefficients when the outcome is regressed on genetic instruments and
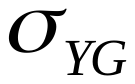
 to be the corresponding standard errors (SE), and
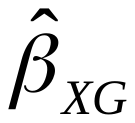
 to be the estimated coefficients when the risk factor is regressed on the genetic instruments with SE
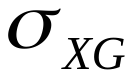
. We also assume the correlation between two genetic variants G_1_ and G_2_ to be
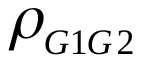
, and
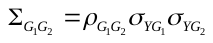
.

The estimate from a weighted generalized linear regression can be formulated by

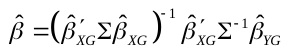

with SE

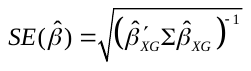

A similar approach may be used for MR-Egger, which allows an intercept term in the weighted regression. Please also refer to ref. ^2^^,^ ^3^ for details. The presence of imbalanced horizontal pleiotropy could be assessed by the significance of the intercept term (testing whether intercept=0). We employed the R package “MendelianRandomization” for the above analyses.

As remarked by Burgess et al.^1^, inclusion of a larger number of variants in partial LD may improve the power of MR, as the variance explained by the instruments might go up. Including “redundant” SNPs in addition to the causal variant(s) will not improve power; on the other hand it will not invalidate the results. However, including too many variants with high correlations may also result in instability of MR estimates^4^. In this study, we performed LD-clumping of genetic instruments at an r^2^ threshold of 0.1, balancing the benefits and risks of using a too high or too low threshold.

References:

1.	Burgess S, Dudbridge F, Thompson SG. Combining information on multiple instrumental variables in Mendelian randomization: comparison of allele score and summarized data methods. *Stat Med* 2016; **35**(11)**:** 1880-1906.

2.	Bowden J, Smith GD, Burgess S. Mendelian randomization with invalid instruments: effect estimation and bias detection through Egger regression. *Int J Epidemiol* 2015; **44**(2)**:** 512-525.

3.	Schmidt AF, Dudbridge F. Mendelian randomization with Egger pleiotropy correction and weakly informative Bayesian priors. *Int J Epidemiol* 2017.

4.	Burgess S, Zuber V, Valdes-Marquez E, Sun BB, Hopewell JC. Mendelian randomization with fine-mapped genetic data: Choosing from large numbers of correlated instrumental variables. *Genet Epidemiol* 2017; **41**(8)**:** 714-725.
