## Supplementary figures and images for "Uncovering bi-directional causal relationships between plasma proteins and psychiatric disorders: A proteome-wide study and directed network analysis"

### Figure S1

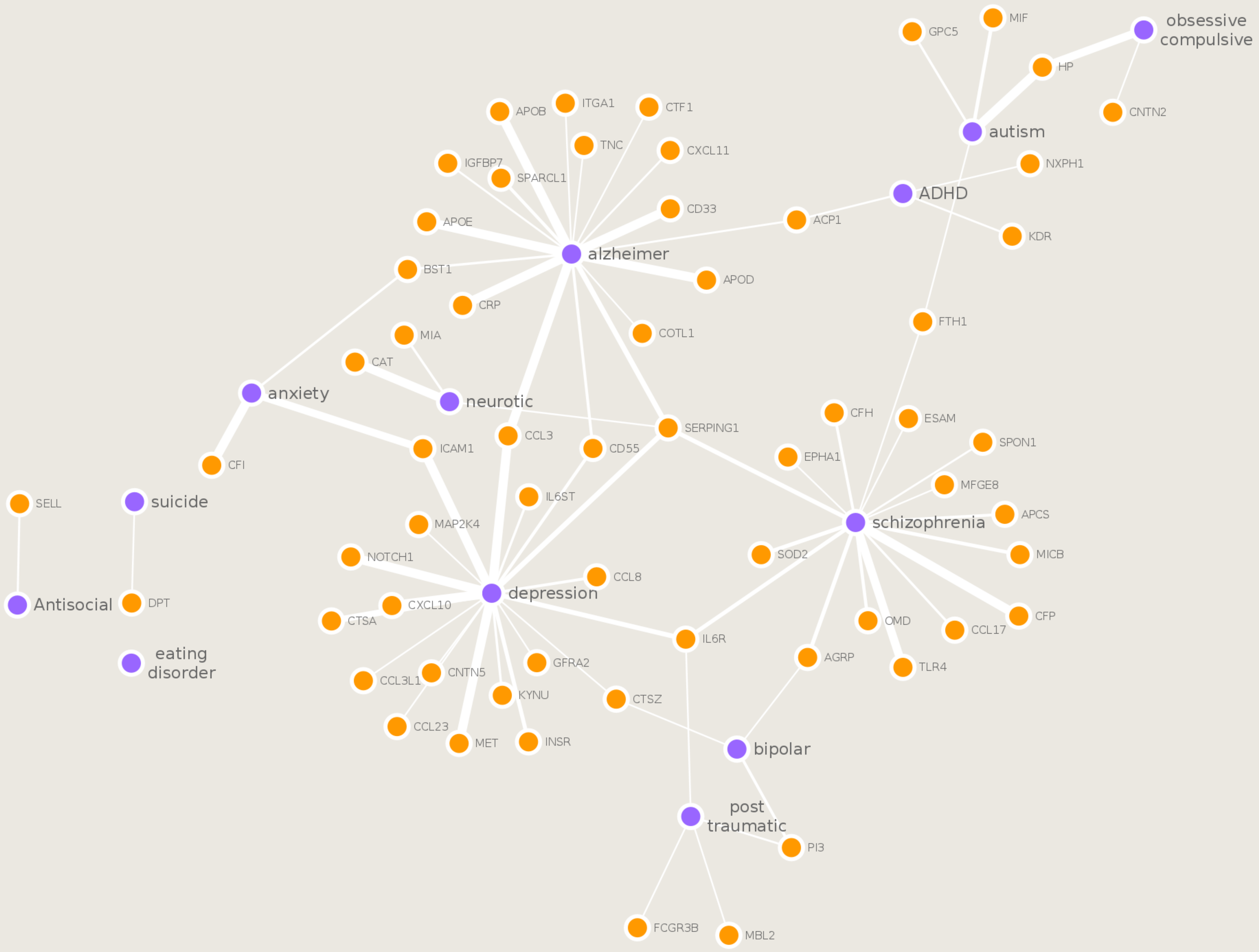

### Figure S2

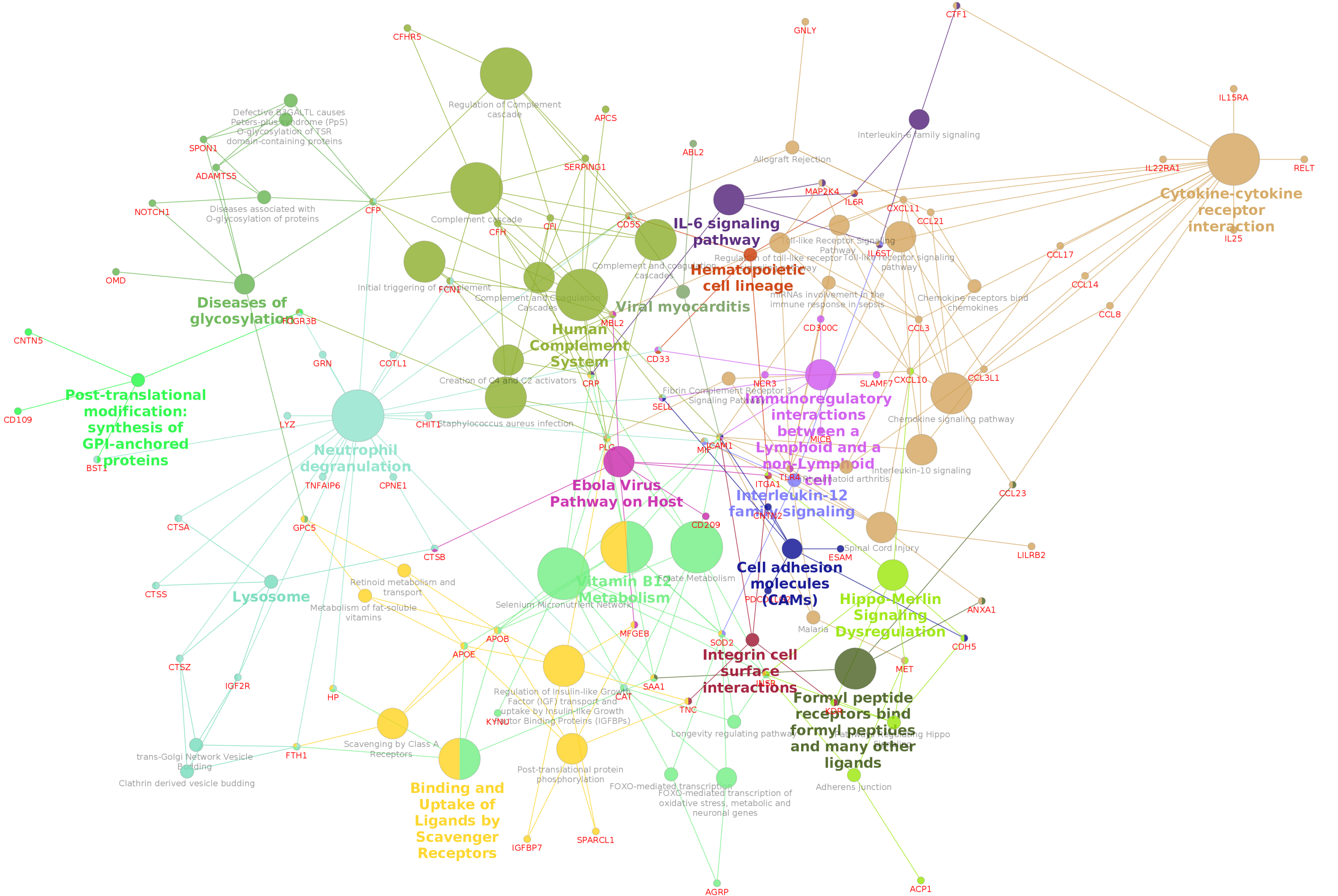
